# Covalent Stabilization of Collagen Mimetic Triple Helices and Assemblies by Dopa Crosslinking

**DOI:** 10.1101/2024.09.30.615841

**Authors:** Carson C. Cole, Brett H. Pogostin, Vardan H. Vardanyan, Kiana A. Cahue, Thi H. Bui, Adam C. Farsheed, Joseph W.R. Swain, Jonathan Makhoul, Marija Dubackic, Peter Holmqvist, Ulf Olsson, Anatoly B. Kolomeisky, Kevin J. McHugh, Jeffrey D. Hartgerink

**Affiliations:** Department of Chemistry, Rice University 6100 Main Street, Houston, Texas 77005, United States; Department of Bioengineering, Rice University 6100 Main Street, Houston, Texas 77005, United States; Division of Physical Chemistry, Lund University, P.O. Box 124, SE-22100 Lund, Sweden; Department of Chemical and Biomolecular Engineering, Rice University, Houston, Texas 77005, USA; Department of Physics and Astronomy, Rice University, Houston, Texas 77005, USA

## Abstract

Creating thermally stable collagen mimetic peptides (CMPs) is a persistent challenge. Nature leverages covalent crosslinkings to stabilize collagen’s signature triple helical tertiary structure and higher-order assemblies. Herein, we demonstrate that crosslinkings between levodopa (Dopa) and lysine, amino acids present in native collagen, can covalently stabilize the triple helix in collagen mimetic peptides. Since alkaline conditions catalyze the oxidation of the catechol on Dopa to a benzoquinone, while being in proximity to the nucleophilic lysine, we hypothesized that this reaction could be a facile method to covalently capture the supramolecular structure of CMPs by simply increasing the pH of the aqueous solvent with the addition of sodium hydroxide. This covalent capture strategy successfully stabilizes CMP homotrimers and a de novo designed ABC-type heterotrimer demonstrating that the Lysine-Dopa covalent bond is best templated by a supramolecular, axial cation–*π* pairwise interaction. In nature, collagen can hierarchically assemble into fibers. This behavior can be mimicked with the self-assembly of CMPs, but the resulting nanofibers typically exhibit thermal stability below body temperature. In a final application, we demonstrate that Dopa–Lysine covalent capture also enhances the thermal stability of CMP nanofibers well above 37 °C. This biomimetic covalent capture strategy can stabilize a wide variety of CMP systems and potentially enable the biomedical application of these materials.

## 1 Introduction

Nature utilizes post-translational modifications (PTMs) to modify protein structures and to direct biological processes. ^1,2^ Post-translational modifications in the proteome include phosphorylation, glycosylation, farnesylation, hydroxylation and many others. ^3^ These crucial chemical modifications occur on both globular and fibrous proteins. The fibrous proteins of the extracellular matrix (ECM) rely on PTMs to guide structure, assembly, and function. ^4,5^ Collagen, the most abundant protein in the ECM, relies heavily on PTMs to guide folding and maintain its correct conformation both in vitro and in vivo. ^6^

A crucial PTM in collagen synthesis in vitro and in vivo includes hydroxylation. Hydroxylation of proline at the 4^*th*^ position is essential in providing stability to collagen’s triple helical backbone by stabilizing the imino ring’s *exo* pucker. ^7,8^ PTMs essential to collagen assembly are also able to facilitate covalent crosslinking. Collagen is a relatively unstable molecule at body temperature. ^9^ To overcome this inherent low stability, collagen crosslinks with itself and other ECM proteins. Active enzymatic crosslinking can involve lysyl oxidases (LOX) and the class of lysyl-oxidase likes (LOXLs), which leads to the oxidative deamination of a *ζ* -amine of Lys to the aldehyde-containing allylysine that can condense with each other or other Lys and hydroxylysine residues. ^10^ There are also a variety of less-known interactions, such as sulfilimines, which are important covalent crosslinks for collagen type IV. ^11^

Recently, it has been reported that collagen type I, the most abundant form of fibrous collagens, contains levodopa (Dopa) residues which are implicated in the scavenging of oxidative radicals. ^12,13^ Dopa, a hydroxylated tyrosine, is known to readily undergo oxidation to a benzoquinone under neutral and basic conditions. ^14^ These reactive quinone species can form Dopa–Dopa covalent crosslinks or react with nucleophiles, such as primary amines, to form Schiff Bases of Michael Addition products (Figure 1a). Covalent bonds between Dopa and Lys are observed in proteins, including the collagen-related enzymes LOX and LOXL, which possess a lysl tyroslquinone (LTQ) crosslinking between a Dopa and Lys residue in the enzyme’s active site. ^15–17^ Considering that natural collagen contains both Dopa and Lys residues, we utilized self-assembling collagen mimetic peptides to study the potential supramolecular and covalent interactions between Dopa and Lys residues in the triple helix.

**Fig. 1.**
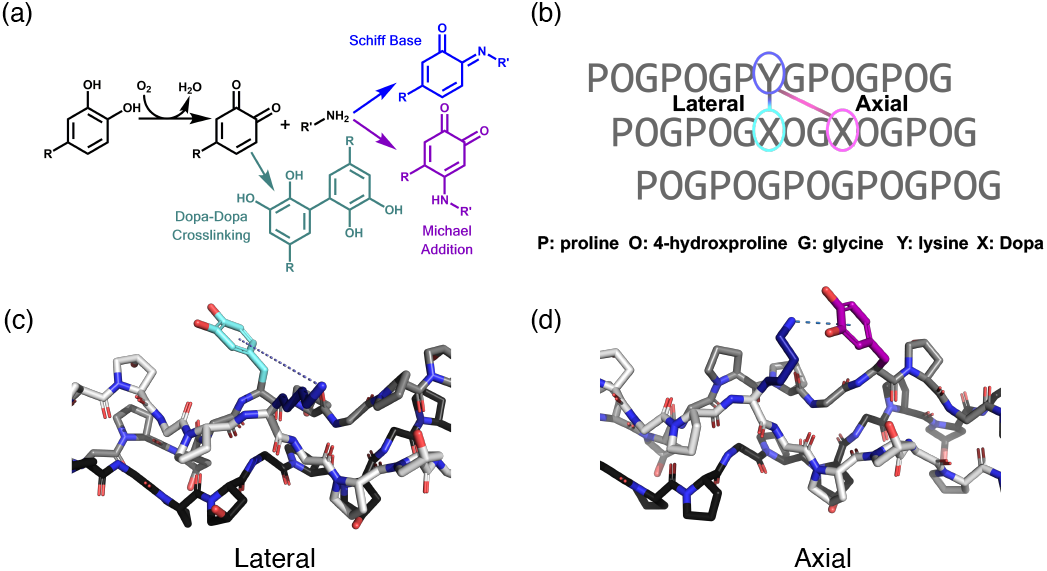
Pairwise interaction geometry and oxidation of levodopa (Dopa) to *o*-benzoquinone. (a) A schematic showing Dopa oxidation and then the subsequent possible Schiff Base and Michael Addition reactions with a primary amine. Dopa-Dopa crosslinking is also a possible product. (b) The prototypical collagen sequence necessary for different interaction geometries with lateral occurring between Yaa (blue) and Xaa of the same triplet (teal) or axial occurring between Yaa (blue) and Xaa of the subsequent triplet (pink). (c) A lateral presentation of Lys (blue) and Dopa (teal). (d) An axial presentation of Lys (blue) and Dopa (magenta). Amino acids were substituted from PDB ID: 3B0S. ^18^.

Self-assembling peptides assemble into a variety of biomimetic nanostructures, including fibers, tubes, and sheets. ^19–21^ Collagen mimetic peptides undergo self-assembly into the right-handed triple helical tertiary structure of natural collagens. ^22^ Collagens have the canonical repetitive amino acid motif of Xaa-Yaa-Glycine, where the Xaa is typically proline, and Yaa is 4-hydroxyproline. The abundance of imino acids leads to rigidity in the three left-handed polyproline II (PPII) that wind into a right-handed superhelix with a periodic symmetry. ^6,18^ The glycine is sterically non-imposing and facilitates an inter-strand hydrogen bonding network between its amide protons and the carbonyl of the Xaa of the adjacent strand. This leads to a single amino acid offset and defines the register of the triple helix between distance leading, middle and trailing strands. ^23^ When the amino acid sequences of all three strands are equivalent, the triple helices are referred to as homotrimers, while triple helices containing nonequivalent strands can be either A2B or ABC heterotrimers. ^24,25^

The winding of the triple helix leads to distinct presentations of the amino acid side chains in a pairwise manner. These pair-wise interactions have two known geometries: axial and lateral (Figures 1b). In most cases, the axial is more geometrically accessible than the lateral and has been used extensively in collagen mimetic peptide design. ^26^ It is also known from both molecular dynamics and experimental evidence that charge pairs, ^27^ amide-*π* ^28^ and cation-*π* ^29,30^ pairwise interactions can effectively stabilize the triple helix. By achieving close proximity, pairwise interactions can also undergo various chemical modifications to create a synthetic, interstrand covalent bond. ^31^ For example, the close interaction distance between residues in a charge-pair interaction allows for proximity-directed amidation and isopeptide bond formation, typically between lysine and glutamate, ^32^ which enhances the thermal stability of the PPII helix through covalent capture ^33^ akin to disulfide bonds. ^34,35^ However, this synthesis requires the use of activating reagents, like hydroxybenzotriazole (HOBt) and 1-ethyl-3-(3-dimethylaminopropyl)carbodiimide (EDC) and has been reported with modest yields. ^36^

To facilitate a reagent-free covalent capture of triple helices, we hypothesized that Dopa could be used to form a bond, similar to the LTQ covalent crosslinking, with the *ζ* -amine of a proximal Lys. ^37,38^ In this work, we investigate collagen triple helix stabilization through the supramolecular cation-*π* pairwise interaction and subsequent reagent-free covalent capture. First, using circular dichroism, we probe the triple helices’ thermal stability for axial and lateral substitution geometries. Then, using molecular dynamics (MD) simulations, we investigate the propensity for the geometry of the triple helix to facilitate either an axial or lateral interaction (Figures 1c/d). By employing an axial Lys–Dopa interaction in a complex ABC-type heterotrimer, we demonstrate that this covalent methodology can be useful in diverse chemical environments with the caveat that supramolecular assembly is necessary to facilitate proximity-directed reactivity. Finally, we show that the cation-*π* interaction can help create a supramolecular collagen mimetic hydrogel and subsequent covalent capture can be used to significantly improve the thermal stability of CMP fibers. This work demonstrates the promise of this reagent-free, Dopa-based covalent capture approach.

## 2 Materials and Methods

### 2.1 Peptide Synthesis

Peptides were synthesized on a solid phase low-loading rink amide MBHA resin (0.35 mmol/gram) using standard Fmoc-protected amino acids to obtain peptides with C-termini amidation. For F*Dopa*, the peptide was synthesized on preloaded Gly-Wang resin, resulting in a free C-terminus. For amino acid deprotection, 25% (v/v) piperidine in dimethylformamide (DMF) was used. Coupling steps were performed at room temperature (ca. 25 °C) with 2-(1H-7-Azabenzotriazol-1-yl)-1,1,3,3-tetramethyl uronium hexafluorophosphate, methanaminium (HATU) and diisopropylethylamine (DiEA) in DMF at 1:4:4:6 equivalents of resin:amino acids:HATU:DiEA. Acetylation of the N-terminus was done using a double addition of excess acetic anhydride and DiEA in dichloromethane (DCM). Peptides were cleaved from the resin with 7.5% v/v scavengers (triisopropylsilane, H_2_O, ethanedithiol and anisole) in trifluoroacetic acid (TFA). While F*Dopa* was not acetylated, resulting in an unprotected N-terminus after TFA cleavage. The TFA mixture was evaporated under a stream of nitrogen gas, and cold diethyl ether was used to precipitate the crude peptide. The crude peptide was centrifuged, the ether was decanted, and the wash was repeated. Peptides were dissolved in H_2_O and filtered for purification. The peptides were purified via reverse-phase high-performance liquid chromatography prep scale (HPLC), with the binary solvent system being water and acetonitrile, each with 0.05% TFA, at a gradient of 0.7%/min on a 19 x 250 mm XC-18 column with a flow of 20 mL/min. The ACN was removed in vacuo and then frozen before lyophilization. Liquid chromatography on an Agilent Pursuit 5 Diphenyl 150 x 2.0 mm column with tandem mass spectrometry (LC-MS) confirmed the peptide mass and purity (Agilent, California, USA). LC and MS for all peptides can be found in SI Figures S1-S8. Pure, lyophilized peptides were prepared at a working concentration in their respective buffers. All peptide concentrations were determined by mass.

### 2.2 Covalent Capture

To prepare peptides for covalent capture, all peptides were dissolved at pH 4.5 and annealed at 85 °C for 15 mins. Once the peptide solutions cooled to ambient temperature, the samples were left to fold for 24 hrs at 4 °C. All covalent capture reactions were performed at a peptide concentration of 3 mM at pH 9.5 for a minimum of 3 days. Buffer and temperature reaction conditions varied as needed between peptide systems to ensure the non-crosslinked triple helix remained stable (supporting information Table S1). For homotrimer reactions, peptides were dissolved at 6 mM in water at pH 4.5 (pH adjustments were made with 1 M HCl/NaOH), annealed, and allowed to fold for at least 24 hours. After a polyproline type II (PPII) signal was confirmed with circular dichroism with a positive peak at 225 nm, the sample was mixed in a 1:1 volume ratio with 20 mM bicarbonate buffer (pH 9.5) to initiate the covalent capture reaction. The pH was adjusted with small volumes of 1M NaOH to ensure the pH remained at 9.5 post-dilution. For heterotrimer reactions, the folded ABC system in 10 mM phosphate buffer dissolved at a total peptide concentration of 3 mM (1mM of each constituent peptide of the heterotrimer). Heterotrimer samples were adjusted to pH 7.4 with 1M NaOH to initiate the reaction. Covalent capture of fiber-forming CMPs (F*Dopa*) was performed in water without any additional buffer, and the pH was adjusted to 9.5 by adding 1M NaOH to initiate crosslinking. The reactions were characterized by matrix-assisted laser desorption ionization (MALDI) TOF MS with a 1% (w/v) matrix of *α*-cyano-4-hydroxycinnamic acid (CHCA) in 50:50 acetonitrile:H_2_O with 0.05 % TFA or by UV-Vis (see SI Methods).

### 2.3 Circular Dichroism

All circular dichroism (CD) data were collected on a J-810 spectropolarimeter (Jasco, Tokyo, Japan) equipped with a Peltier temperature controller. Spectrum measurement was collected at 5 °C. Heterotrimer and homotrimer samples were diluted into a 1 mm cuvette with fresh MQ H_2_O for a final peptide concentration of 0.3 mM and 1 mM buffer with a pH = 4.5. The melting curves were collected from 5 °C to 85 °C with a heating rate of 10 °C/hour. To monitor refolding, the sample solution was kept at 85 °C for 30 min, and then the reverse temperature gradient was applied back to 5 °C. The first-order derivatives of the melting curves were calculated with the Savitsky-Golay smoothing algorithm, and the temperature where the minimum derivative value appears was defined as the melting temperature (*Tm*). ^39^ The mean residue ellipticity (MRE) value was then calculated with the equation, MRE = (*θ* × m)/ (c × l × nr × 10) where *θ* represents the experimental ellipticity in millidegrees (mDeg), m is the molecular weight of the peptide (g/mol), c is the peptide concentration (milligrams/milliliter), l is the path length of the cuvette (cm) and nr is the number of amino acid residues in the peptide.

### 2.4 Molecular Dynamics Simulations

All-atom molecular dynamics (MD) simulations were performed to estimate the distance between the Lys side chain’s nitrogen and Dopa’s benzene ring. The initial collagen structure was obtained from the Protein Data Bank (PDB ID: 3B0S). ^18^ By using DLPacker, ^40^ a molecular packing software, energy-minimized R-groups for both Lys and Dopa were incorporated into the structure at the 12^*th*^ and 14^*th*^ positions. The simulation was prepared using the CHARMM-GUI. ^41,42^ The cubic simulation box had a 12 nm side and approximately 118,000 atoms. The solution was comprised of 39198 TIP3P water molecules and 150 mM NaCl counter-ions. Gromacs ^43–45^ was used to carry out MD simulations and was parameterized with a Charmm36 force field. ^46^ The simulations employed periodic boundary conditions in all directions, and electrostatic interactions were treated by using the particle-mesh Ewald method. ^47,48^ The simulation system was minimized and thermally equilibrated under an NVT ensemble. After thermal equilibration, we carried out pressure equilibration under an NPT ensemble. Equilibration was performed using a 1 fs simulation step. After equilibration, we increased the simulation step to 2 fs and ran the production MD for 150 ns. The coordinates of the simulation system were extracted every 100 ps. Visual Molecular Dynamics (VMD) ^49^ software was used to analyze the simulation data.

### 2.5 NMR

The isotopically-labeled ABC-type heterotrimer peptides were prepared at 2.7 mM in 9 mM phosphate buffer, 10 mM trimethylsilyl propanoic acid (TSP), and 90%H_2_O/10%D_2_O. All characterizations were performed on a Bruker NEO 600 MHz High-Performance digital NMR with a helium-cooled inverse TCI probe at 30 °C. All ^1^H-^15^N HSQC data was collected using the same set of parameters, including 1024 x 128 complex points, 8 scans, and 16 dummy scans. A spectral window of 35 ppm was used for the nitrogen dimension and 16 ppm for the hydrogen dimension. The nitrogen carrier frequency was set at 117 ppm, and the proton carrier frequency was set to match the water signal. The 90-degree pulse was calibrated separately for each sample. Raw NMR data was processed using TopSpin 4.3.0 software (Bruker, Mass., USA).

### 2.6 Cryo Electron Microscopy

F*Dopa* was prepared at 2.00% w/v in MQ H2O (pH 4.5 and 9.5). The pH was adjusted to 9.5 by adding 1M NaOH to initiate crosslinking. Before plunging, samples were diluted to 0.10% w/v in fresh MQ H_2_O. Samples were prepared on 200 mesh lacey carbon grids. 3 *µ*L of each sample were plunged into liquid ethane with a Leica EM GP automatic plunge freezer (Leica Microsystems, Wetzlar, Germany). The plunging chamber was set to have the following conditions: a temperature of 20 °C and a humidity of 90%. After preparation, samples were examined at the National Center for High-Resolution Electron Microscopy on a JEM-2200FS (JOEL, Tokyo, Japan) transmission electron microscopy instrument using an F416.0 camera (TVIPS, Gauting, Germany) with an accelerator voltage of 200 kV. Zero-loss images were acquired using Serial EM in a low-dose mode.

### 2.7 Scanning Electron Microscopy

F*Dopa* was prepared at 2.00% w/v in MQ H2O (pH 4.5 and 9.5). The pH was adjusted to 9.5 by adding 1M NaOH to initiate crosslinking. Samples were placed into Porous Spec Pots (Electron Microscopy Sciences, Hatfield, PA) and were dehydrated using a series of ethanol in MQ H_2_O in a series of dilutions (30, 50, 60,70, 80, 90%, 2 × 100%) for 10 min each. The samples were then critical point dried using a Leica EM CPD300 (Leica Biosystems, Deer Park, IL), followed by sputter coating with 5 µm of gold using a Denton Desk V Sputter System (Denton Vacuum, Moorestown, NJ). SEM was done using a Helios NanoLab 660 Scanning Electron Microscope (FEI Company, Hillsboro, OR) and all micrographs were obtained at 2 kV and 25 pA.

### 2.8 Scattering

Small-angle X-ray scattering measurements were performed using the laboratory-based Xeuss 3.0 instrument (Xenocs, France) at The Center for Scattering Methods (CSM) in the Faculty of Science, Lund University, equipped with an X-ray source producing a photon beam with a wavelength of 1.34 Å. The scattering patterns were recorded with an Eiger2 R1M 2D-detector (Dectris) and azimuthally integrated using the XSACT program available with the equipment, creating 1D scattering curves. The radially averaged intensity, I(q), is given as a function of the scattering vector *q*=4*π* sin*θ* /*λ* , where *λ* is the wavelength and 2*θ* is the scattering angle. F*Dopa* was prepared at 2.00% w/v in MQ H2O (pH 4.5 and 9.5). the pH was adjusted to 9.5 by adding 1M NaOH to initiate crosslinking. The sample capillaries were put in a Peltier-type multi-sample holder, heating the samples from 5 to 60 °C in 5 °C steps. Samples were equilibrated for a minimum of 10 minutes at each temperature before scattering measurements were taken. The fitted scattering curves at 5 °C were fit using SasView 5.0.5 (fitting parameters shown in Table S2).

## 3 Results and Discussion

### 3.1 Covalent Capture of Collagen Homotrimers

To investigate if crosslinking between Lys and Dopa residues could stabilize a triple helix, we synthesized two peptides: OGDopaK and KGDopaO. Two control peptides were also synthesized, termed AGDopaO and KGDopaO6,9*GlyDeletion*, to investigate the propensity for side reactions, including the self-reactivity of the *o*-benzoquinone (Table 1). In OGDopaK, leading to middle (L-to-M) and middle to trailing (M-to-T) strands’ interactions between Lys and Dopa are both in the lateral geometry. Meanwhile, the trailing to leading (T-to-L) strands are too distant and expected to be non-interacting. The substitution scheme of KGDopaO presents the Lys and Dopa with an axial interaction in the L-to-M and M-to-T, while the T-to-L is a lateral interaction. ^50^ After synthesis and folding at 3 mM (pH 4.5), we observed that all three homotrimers formed with a positive PPII signal at 225 nm as observed with circular dichroism (CD) (Figure S9).

**Table 1.**
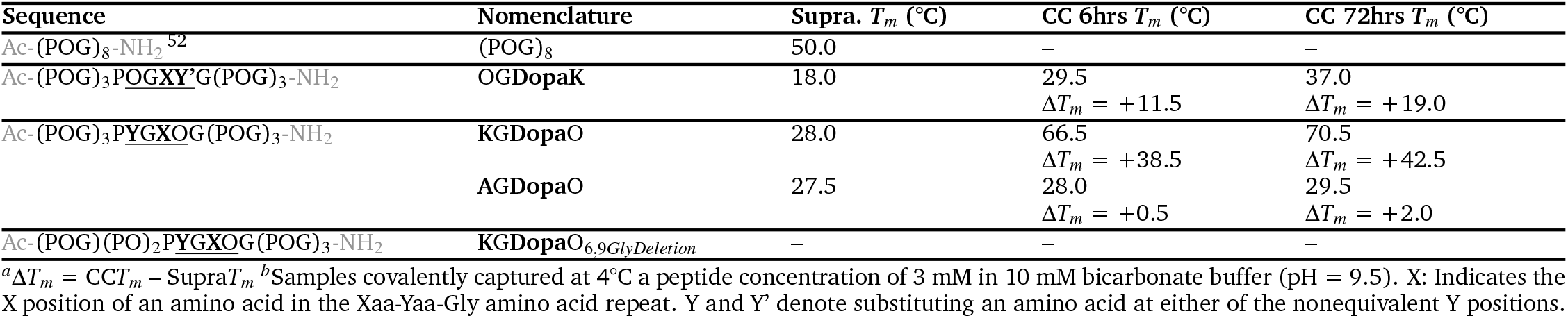
Enhanced Stabilization with Covalent Capture (CC)

By monitoring the CD thermal unfolding curves, we were able to measure the covalent stabilization of the triple helix by oxidation induced by a switch to pH 9.5 and subsequent covalent bond formation between the Lys and Dopa. The oxidation of Dopa to an *o*-benzoquinone can be accomplished at basic pH while simultaneously enhancing the nucleophilicity of the *ζ* -amine of Lys (lysine-N*ζ* , pK*a* ≈ 9.3 – 13.2^51^). The thermal transition temperature (*Tm*) is defined by the minimum of the first derivative of the thermal unfolding curve. The supramolecular and covalently captured triple helix *Tm* values are summarized in Table 1. As in previous studies, the substitution of the pairwise cation-*/pi* interactions lowered the thermal stability of the supramolecular assembly when compared with a peptide (POG)_8_ with only proline and hydroxyproline in the X and Y amino acid positions of the Xaa-Yaa-Gly repeat. ^52^

The presentation of Lys – Dopa interaction in the lateral geometry is accomplished in the OGDopaK homotrimer. There was a steady increase in thermal stability (Figure 2a). Refolding curves in Figure 2b show that the covalently captured homotrimer after 72 hrs had the highest recovery. However, the asymmetry of the first-order derivative melt curves in Figure 2c suggests heterogeneity in the covalently captured samples. This decreased cooperativity is also supported by the MALDI-MS data in supporting information Figure S10b, showed peaks correlating to dimer, trimer, tetramer and possibly higher-order species. Nevertheless, there was a 19.0 °C stabilization for OGDopaK when compared with the non-crosslinked triple helix.

**Fig. 2.**
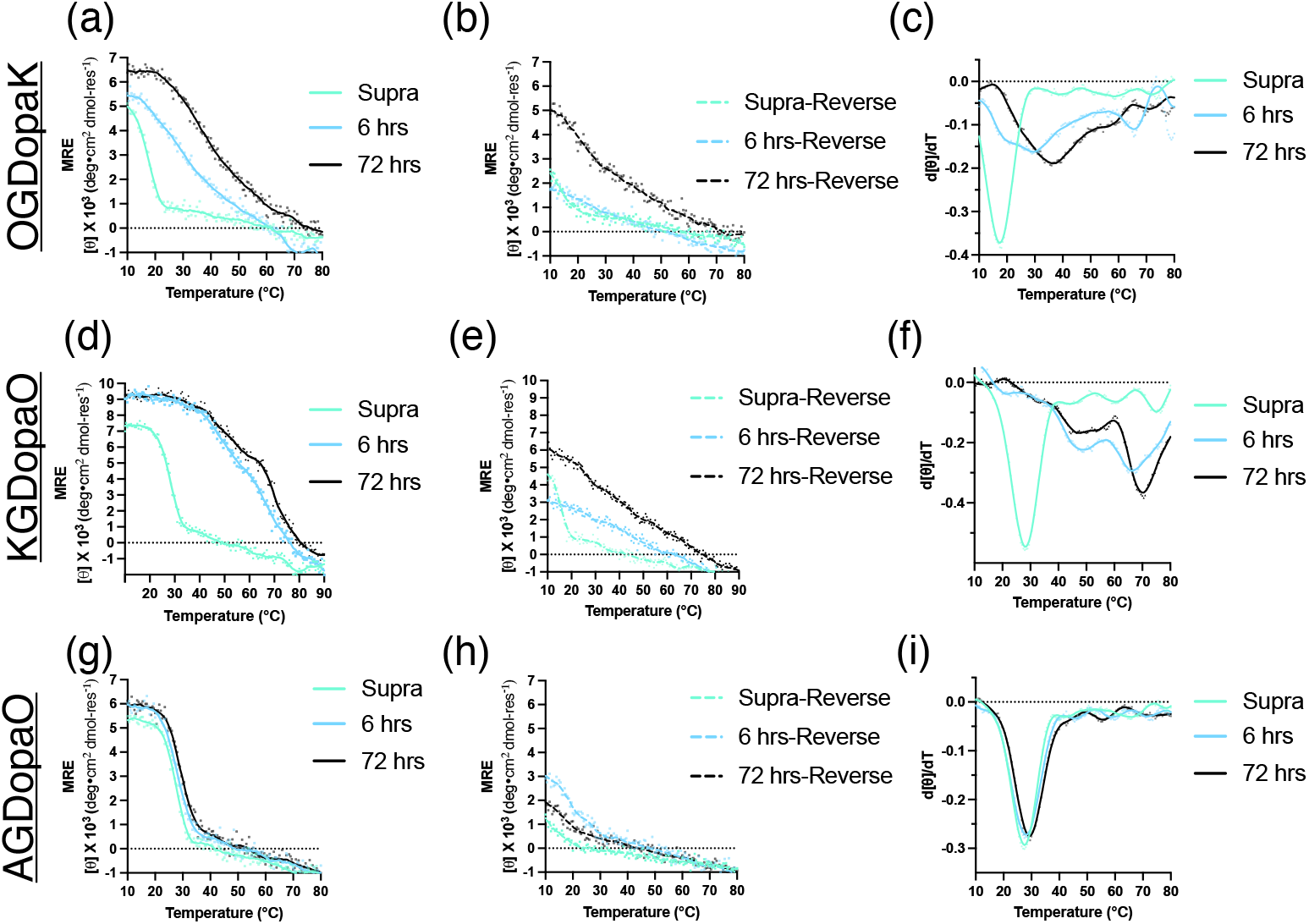
Reaction kinetics of OGDopaK, KGDopaO and AGDopaO homotrimers at basic pH monitored by circular dichroism. (a) CD thermal melt curves for OGDopaK from the supramolecular assembly to 72 hrs of covalent capture at pH 9.5. (b) CD thermal reverse melts (dashed lines) demonstrate a change in hysteresis (c) First-order derivatives of OGDopaK containing peptides, where the minima denote the transition temperature (*T*_*m*_). (d) CD thermal melt curves for KGDopaO from supramolecular to 72 hrs. (e) CD thermal reverse melts (dashed lines) demonstrate a change in hysteresis, where MRE recovery is greatest at 72 hrs (f) First-order derivatives of KGDopaO containing peptides, where the minima denote the *T*_*m*_ and stabilization of 48.5 °C at 72 hrs. (g) CD thermal melt curves for AGDopaO from supramolecular to 72 hrs. (h) CD thermal reverse melts (dashed lines) demonstrate a little to no change in hysteresis (i) First-order derivatives of AGDopaO containing peptides, where the minima denote the *T*_*m*_. For CD, raw data are presented as points, and lines are a 3rd-order Savitsky-Golay fit. ^39^

Next, we investigated the propensity of stabilization for an axial interaction in the homotrimer KGDopaO. The cation-*π* supramolecular interaction and subsequent covalent capture is favored in the axial conformation and supported by the characterization of KGDopaO (Figures 2d-f). The non-crosslinked triple helix had a 10.0 °C difference in stability when compared with OG-DopaK and agrees with previous studies on cation-*π* interactions in triple helices, demonstrating a preference for the axial geometry. ^30^ After covalent capture at pH 9.5, MALDI ToF MS showed dimer, trimer, and tetramer peaks (see supporting information Figure S10a). This supports the lack of specificity and the observed uncontrolled reactivity common in catechol crosslinks. ^53^ We also observed an asymmetry in the first derivative melting curve of the CD spectra, suggesting multiple species. However, the sample does not possess a thermal transition as broad as that of OGDopaK. After 72 hours, the most stable species observed in CD for KGDopaO gained a remarkable 42.5 °C. For comparison, previous interstrand stabilization attempts with cysteine and homocysteine were either destabilizing or negligible ^34^, and isopeptide amide bond formation provided a gain of 43.5 °C. ^33^

Next, we repeated the crosslinking experiment for two additional peptides, AGDopaO and KGDopaO6,9*GlyDeletion*, to further confirm the axial interaction is preferred for covalent stabilization. AGDopaO, which was synthesized to account for possible Dopa-Dopa crosslinking interactions (Figure 1a), showed a trivial gain in stability of 2.0 °C. In addition, there was a lack of change in the hysterses of the reverse melts presented in Figure 2h, suggesting that Dopa-Dopa covalent interactions are negligible in stabilizing the triple helix, despite some amount of dimer formation being observed with MALDI ToF MS (Figure S10c). It was also noticeable that a Lys – Dopa containing peptides exhibited a yellow solution with a corresponding UV-Vis peak at roughly 355 nm (see supporting Figures S11a/b and S13). In contrast, AGDopaO had a UV-Vis peak maximum that red-shifted to 495 nm in Figure S11c, and the solution was visually red, further confirming that KGDopaO and AGDopaO reactions are distinct.

KGDopaO6,9*GlyDeletion* was also used as a control to understand random polymerization events between Lys and Dopa. Glycine, as already stated, is crucial for an interstrand hydrogen bond in natural collagen ^22^ and collagen mimetic peptides. ^54^ Removal of two glycines at positions six and nine prevented triple helix folding, which was not improved upon raising the pH to initiate covalent capture (Figures S14b/c). This supports the idea that the templating of the triple helix, especially in the axial case, is essential for promoting stabilization. This motivated us to further probe the nuances of the supramolecular interaction’s geometry and the distribution of interactions between Lys with various positions in the *o*-benzoquinone to form either a Schiff Base or Michael Addition product.

### 3.2 Cation-*π* Interaction Propensity with Molecular Dynamics

The supramolecular assembly of the collagen triple helix can be used to guide covalent capture of the resulting tertiary structure. To leverage this supramolecular templating for Lys – Dopa covalent capture, understanding how the pairwise cation-*π* interactions between Dopa and Lys influence triple helical stabilization is crucial. Like the covalent bond formation of cysteine, ^35^ homocysteine, ^34^ and charge pair isopeptide bond formation ^36^, the cation-*π* pairwise interaction of Lys – Dopa is mediated, in part, by the distance of their supramolecular interactions. ^55^

By utilizing molecular dynamics simulations, we observed the interaction propensity for both an axial, which occurs from the L-to-M and M-to-T, and the lateral interaction, from the T-to-L strands, in the homotrimer of KGDopaO. The interaction distance between the center of the ring of Dopa and the amine of Lys are correlated in Figure 3a for the axial interaction. With an ideal interaction distance for pairwise cation-*π* interactions being less than 5 Å, we observed that the axial interaction had a significant amount of the population density within this cutoff. In Figure 3d, the lateral interaction shows a marked shift towards a greater distance, which is consistent with previous literature. ^30^

**Fig. 3.**
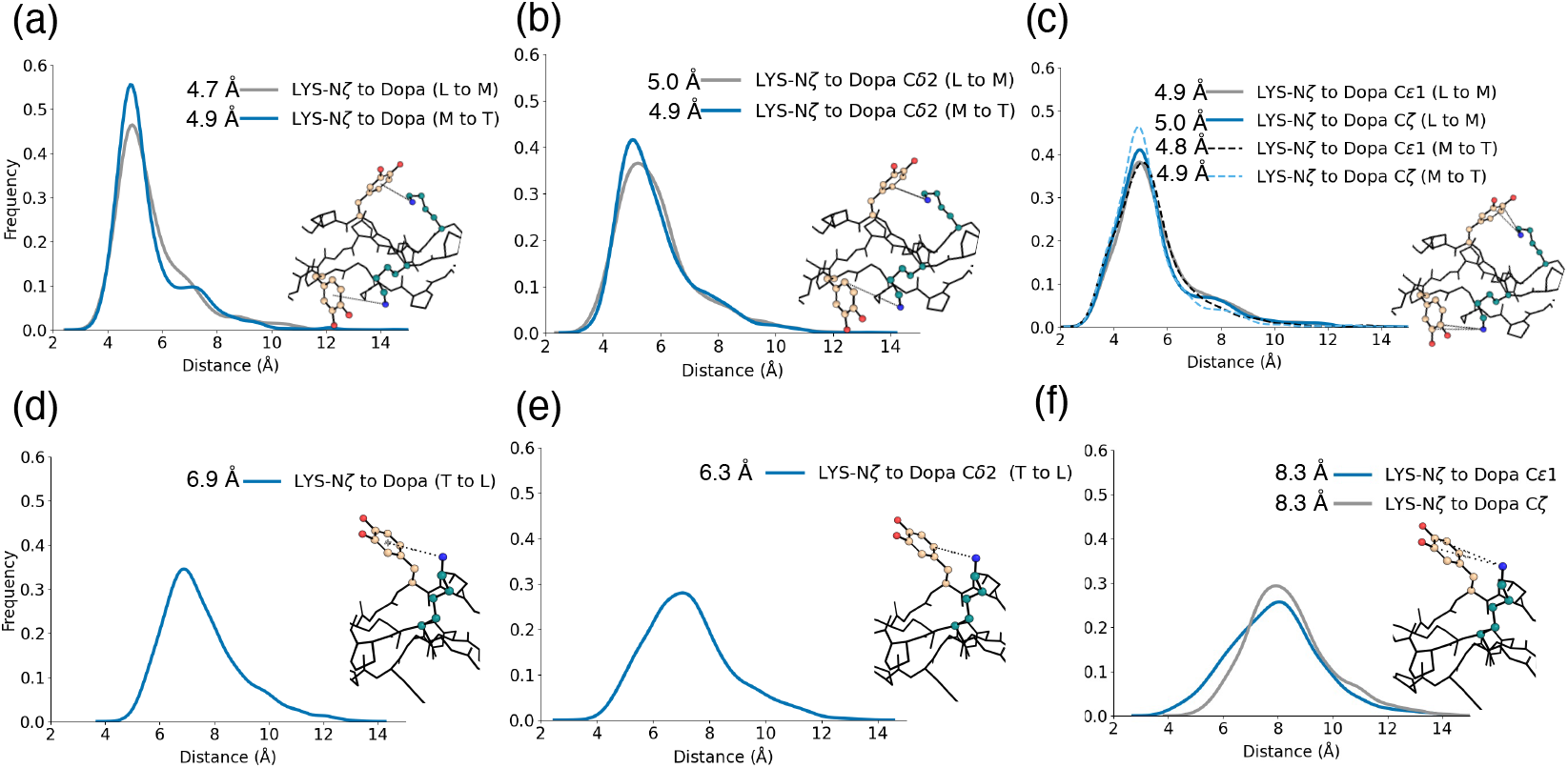
Molecular dynamics plots of distance correlation between Lys and Dopa in axial and lateral interaction geometries within KGDopaO. (a) Plotted frequencies of distances and correlated energy minimized structure between the *ζ* -amine of Lys and the center of the benzyl ring for each pair of interacting amino acids in an axial geometry. (b) Plotted frequencies of distances and correlated energy minimized structure between the Lys *ζ* -amine and Michael-addition site (C*δ* 2) of the benzyl ring for each pair of interacting amino acids in an axial geometry. (c) Plotted frequencies of distances and correlated energy minimized structure between the Lys *ζ* -amine and Schiff Base sites (C*ε*1 and C*ζ* ) of the benzyl ring for each pair of interacting amino acids in an axial geometry. (d) Plotted frequencies of distances and correlated energy minimized structure between the Lys *ζ* -amine and the center of the benzyl ring for each pair of interacting amino acids in a lateral geometry. (e) Plotted frequencies of distances and correlated energy minimized structure between the Lys *ζ* -amine and Michael-addition site (C*δ* 2) of the benzyl ring for each pair of interacting amino acids in a lateral geometry. (f) Plotted frequencies of distances and correlated energy minimized structure between the Lys *ζ* -amine and Schiff Base sites (C*ε*1 and C*ζ* ) of the benzyl ring for each pair of interacting amino acids in an axial geometry. Maxima of each distribution are denoted next to each line, where axial interactions are generally in closer proximity than lateral interactions.

Covalent bonding between Lys and Dopa is possible through Michael Addition or the formation of a Schiff Base (Figure 1a). Thus, we correlated the distance between the nucleophilic amine of Lys and each carbon of Dopa. The Michael Addition product is possible through the reaction of the Lys primary amine with the C*δ* 2 position of the ring (Figures 3b/e). Once again, we observed that the axial interactions would have a greater likelihood of being within 5 Å of this addition site. We also measured the angle (*θ* ) between the normal of the center of the aromatic ring and the Lys *ζ* -amine for the cation-*π* interaction. Where 0° and 180° represent the normal vector, and 90° is correlated to the edge of the aromatic ring. In lateral cases, the angle was closer to 90° with population maxima at 59.58° and 118.69° (see supporting information Figure S15a). Axial interactions were more likely to be located over the ring with maxima from the L-to-M strands having angles of 22.67° and 156.94° and M-to-T strands 22.63° and 157.20° (Figures S15b/c).

The propensity to form a secondary ketimine (Schiff Base) at one of the ketone functional groups was also investigated with molecular dynamics by plotting the normalized frequency between each of the possible addition sites (C*ε*1 or C*ζ* ) of Dopa to the N*ζ* -amine of Lys. In Figures 3c/f, there is again a preference for an axial interaction over a lateral interaction. Compared with the Michael Addition axial plot in Figure 3b, the C*ζ* had higher counts; however, simulations suggest that both the Michael Addition product and Schiff Base can form in this system. These molecular dynamics data indicate that the axial over lateral geometry is more accessible for covalent capture via Dopa oxidation, consistent with the CD thermal stability data in Figure 2.

### 3.3 Covalent Capture of a Complex Heterotrimer

With homotrimer kinetics formally characterized with both experimental and computational techniques, we then incorporated the Lys – Dopa interaction into a complex ABC-type heterotrimer. Covalent capture strategies have proven successful in creating discrete triple helices with greater thermal stability, ^32^ enhanced integrin binding ^56^ and elucidated type I collagen’s heterotrimeric strand registration. ^57^ ABC-type heterotrimers, where the leading, middle and trailing strands are nonequivalent, are particularly challenging to stabilize because of the need for a definite register. ^52,58^ To confirm that we could selectively stabilize an ABC-type triple helix through Lys – Dopa reagent-free covalent capture, we synthesized the ABC-type heterotrimer presented in Table 2. The pairwise interactions promoting heterotrimer formation include axial, lateral charge pairs and axial cation-*π* interactions (Figure 4a). This heterotrimer sequence was based on a previous design and crystal structure (PDB ID: 8TW0) where the amino acid in position 22 of the C strand (C-22), denoted with an X, was previously an aspartic acid (D) (HT3 C - D22). ^59^ The C-22 position was substituted with Dopa to form an axial cation-*π* interaction. Then after covalent capture the axial interaction of B-20-Lys – C-22-Dopa could result in a covalent dimerization from the middle to trailing strands, similar to what Tanrikulu has accomplished with disulfide bridges. ^60^

**Table 2.**
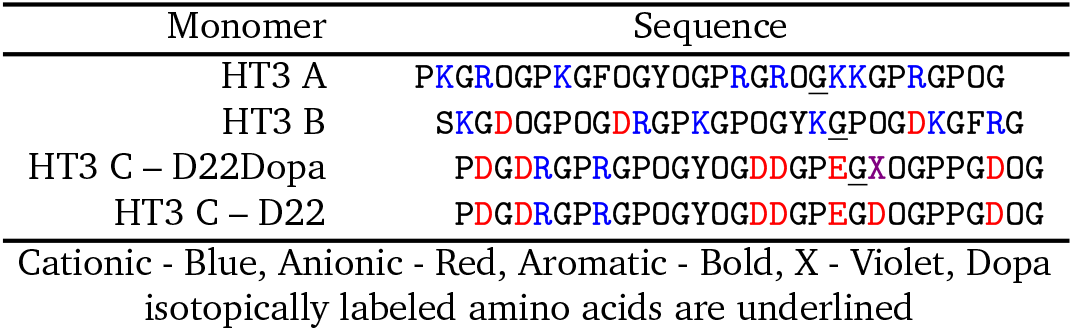
ABC-type Heterotrimer Sequence for Covalent Capture.

**Fig. 4.**
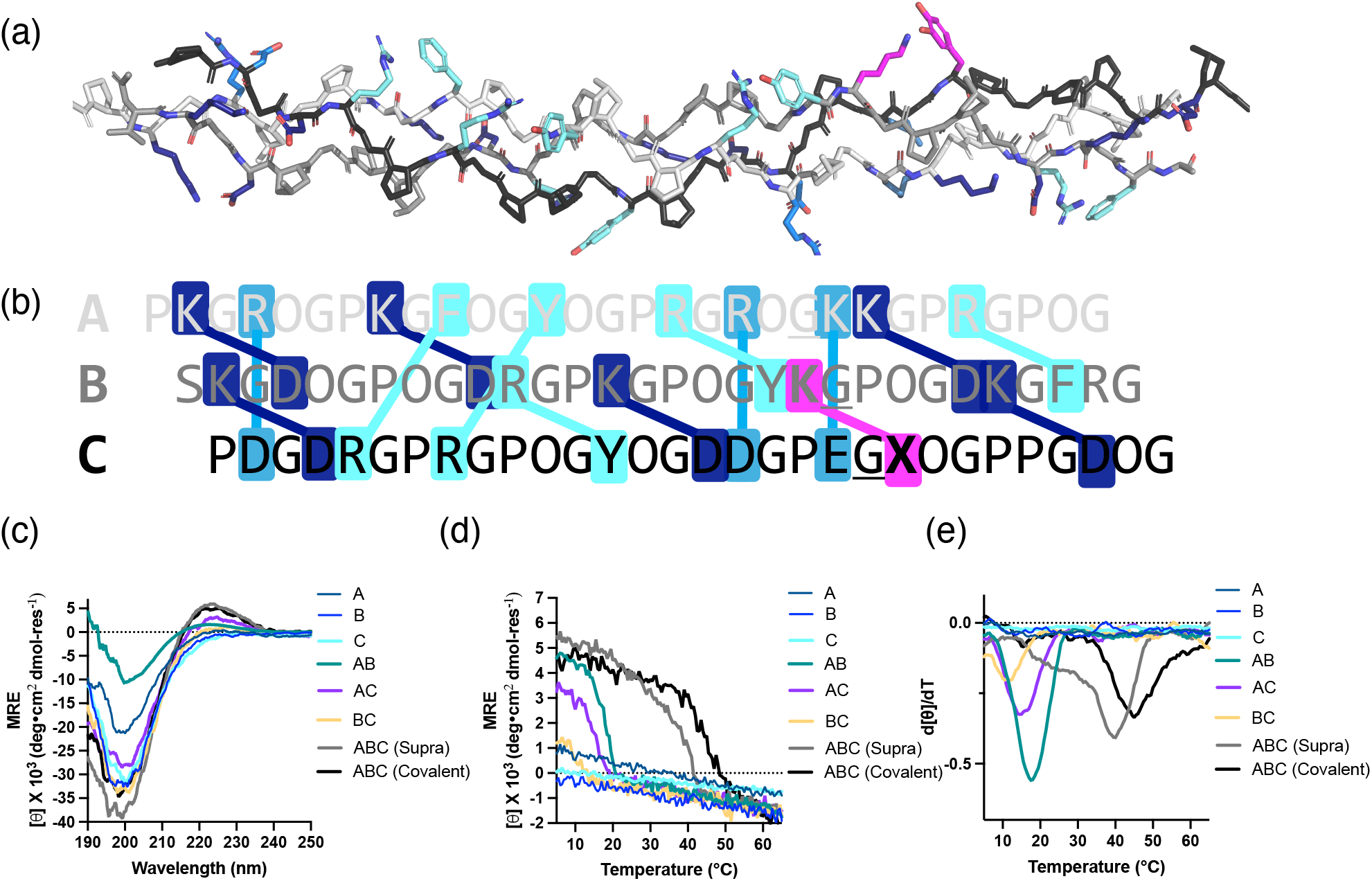
Covalent capture of a complex heterotrimer. (a) A rendering of an ABC-type heterotrimer that is substituted to become ABC – D22Dopa. The leading strand is white, the middle is gray, and the trailing is black. Dark blue amino acid side chains are electrostatic axial, and the lateral interactions are light blue. Teal interactions are cation-*π*, and the magenta pairwise interaction is where the ABC – D22 was substituted to encompass Dopa. Triple helix backbone and non-substituted amino acids are taken from PDB ID: 8TW0. ^59^ (b) Stabilizing pairwise interactions with their relative interaction strength are highlighted in blue, in descending interaction strength: axial charge pairs (K – D), cation-*π* pairs (R – Y) and lateral charge pairs (D – R). In black and purple is the axial Lys – Dopa pairwise interaction. (b) Circular dichroism (CD) spectra at 5 °C of each unary, binary and ternary mixtures. All supra samples are at pH = 4.5, and covalent capture was carried out at pH = 7.4. (c) CD thermal melt curves of all equivalent possible mixtures of the ABC-type heterotrimer at 0.3 mM in 1 mM phosphate buffer. (d) First derivatives of the CD thermal melt unfolding curves.

To validate the unique composition of an ABC-type heterotrimer, we performed CD thermal melts on each ternary, binary and unary mixture. From Table 3, we observed that none of the unary solutions had observable thermal transitions, but the binary mixtures all folded to varying extents. Each binary (AB, AC and BC) and ternary mixture of HT3 ABC ABC - D22Dopa (ABC-Supra and ABC-Covalent) demonstrated a positive CD signal at roughly 225 nm and a negative signal around 198 nm (Figure 4b). In addition, it was evident in both the melt and melt derivative spectra that the ABC, ternary mixture was uniquely stable with a *Tm* = 40.0 °C (Figures. 4b/c). The ABC heterotrimer folded over its second most stable assembly, formed by the mixture of AB components (*Tm* = 17.5 °C) by 22.5 °C, suggesting a unique composition for the ternary mixture. Due to the potential disruption of heterotrimer folding under basic conditions, we covalently captured the folded heterotrimer at pH 7.4. Visually, we observed that the sample turned yellow, suggesting oxidation at neutral conditions. In addition, The covalently captured ABC-type heterotrimer increased the *Tm* by 5.5 °C to a stability of 45.5 °C after 72 hrs (Figure 4d).

**Table 3.** *T*_*m*_ of Binary and Ternary Mixtures for an ABC-type Heterotrimer.

| Mixture | ABC – D22 | ABC – D22Dopa | AB | AC | BC | A | B | C |
| --- | --- | --- | --- | --- | --- | --- | --- | --- |
| Supramolecular | 44.0 °C<sup>a</sup> | 40.0 °C | 17.5 °C | 15.0 °C | 11.5 °C | –<sup>b</sup> | –<sup>b</sup> | –<sup>b</sup> |
| Covalent |  | 45.5 °C |  |  |  |  |  |  |
| $\Delta T_m$ | | +5.5 °C | | | | | | |

<sup>a</sup> Sample was prepared was measured at pH 7.4 as described previously.<sup>59</sup> <sup>b</sup>No strong thermal transition observed Supramolecular spectra were collected at pH = 4.5. CC was performed at 3 mM, pH = 7.4 in 10 mM phosphate buffer and spectrum was collected after 72 hours.

To confirm that the unique ABC composition and register were not compromised by covalent bond formation we performed a 2D (^1^H – ^15^N) heteronuclear sequential quantum coherence (HSQC) NMR study of the ABC-type heterotrimer and compared it against the supramolecular assembly. ^61^ Each strand was labelled with an ^15^N isotopic tag at the 21st glycine (See Table 2). It was hypothesized that the B-20-Lys – C-22-Dopa interaction would alter the chemical shift of the C-21-Gly position and possibly impact the chemical shifts of all tagged amino acids if the triple helix was significantly compromised. When overlaying the HSQC spectra of the ABC supramolecular assembly at pH = 4.5 (Figure 5a), it was observed that there were three peaks correlating to the free monomer strand and three trimer peaks. These trimer peaks (T*x*, where x = A, B or C) closely resemble the shifts of the previously synthesized ABC-type heterotrimer. ^59^

**Fig. 5.**
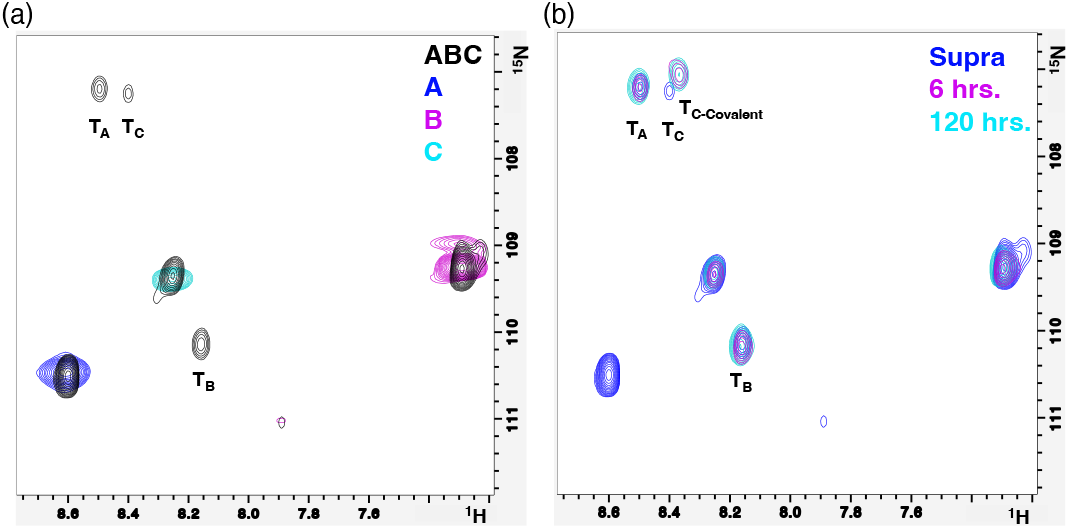
HSQC NMR confirms a unique ABC-composition is maintained upon covalent capture. (a) Overlay of each HSQC of monomers A (dark blue), B (purple) and C (light blue). The supramolecular ABC mixture (black) shows corresponding peaks to each monomer and three distinct trimer peaks (T_*x*_) which can be correlated with their respective strands based upon previous assignment. ^59^ All spectra were acquired at pH = 4.5. (b) The covalently captured sample, adjusted to pH = 7.4, is overlaid with the HSQC spectra of the supramolecular ABC mixture. After 120 hrs, the only observed peak shift is the T_*C*_ signal, which is adjacent to the covalent bond forming with C-22-Dopa. All spectra were acquired at 2.7 mM of total peptide, 10 % D_2_O (v/v) in 9 mM phosphate buffer and at 30 °C.

Next, the pH of the ABC mixture was adjusted to pH = 7.4, and spectra were acquired at six hours to match CD and 120 hours (5 days). Notably, no additional peaks were observed, but the peak corresponding to the C-strand in the ABC-type heterotrimer did shift upfield in the ^1^H and ^15^N dimensions. The B-strand of the trimer (T*B*) also had a minor shift after covalent capture. Both peaks were consistent from the 6 hrs to 120 hrs time points, suggesting that the triple helix is not compromised for up to five days. The T*A* strand’s chemical shifts are constant upon covalent capture. These NMR data confirm that the ABC-type heterotrimer has a unique register and composition, and the ABC-type heterotrimer is maintained after covalent capture.

### 3.4 Covalent Stabilization of Polymerized Collagen Mimetic Fibers

With covalent capture characterized in homotrimeric and heterotrimeric collagen mimetic peptides, we moved to stabilize interhelical interactions with Lys – Dopa covalent capture. Fibrillogenesis of collagen is a complicated assembly process involving different sub-types of collagen, glycosylation, and active enzyme-mediated folding. ^62^ Utilizing supramolecular interactions, such as charge pairs, ^63^ polymerizing collagen mimetic peptides have been shown to form higher-order structures. While these CMPs only employ 33 – 48 amino acid residues compared to the roughly 1,000 residues in natural collagen Type I, biomimetic properties such as D-banding have been conserved. ^64^ These CMP fibers have also been proposed to utilize a sticky-ended ^65^ or symmetric fiber assembly mechanisms. ^66,67^

To further show the utility of this covalent capture method, we substituted Dopa into the sticky-ended polymerizing collagen scheme that contains the sequence of (PKG)4(POG)4(DOG)2XOGDOG (Figures S16a/b). When X = aspartic acid (D), this relatively short polypeptide polymerizes into a nanoscale, fibrous assembly that forms a hydrogel when utilizing only charge pairs. ^63^ When we incorporated the supramolecular cation-*π* interaction, where X = Dopa and termed F*Dopa*, the sample demonstrated pH dependence and formed a hydrogel at pH 4.5. In contrast, at pH 7.4 and 9.0, the immediate oxidation caused precipitate to form (Fig S16c). At pH 4.5, a 2% w/v solution of F*Dopa* in H2O was prepared and formed a hydrogel. The hydrogel had a storage modulus of 640 Pa and loss modulus of 90 Pa and was linearly viscoelastic up to 6.5% strain (Figure S16d). Subjecting the hydrogel to higher strain resulted in a reduction of the storage modulus, indicating that the hydrogel is shear thinning. At a strain of ca. 10%, the loss modulus exceeds the storage modulus, and the material begins to flow like a liquid, losing its hydrogel character.

Using cryo-electron microscopy (Cryo-EM), we were able to visualize the collagen mimetic polymeric mesh of the supramolecular hydrogel at pH 4.5 (Figure 6a). Subsequently, we covalently captured the system by adjusting the hydrogel to pH 9.5. Upon pH adjustment, the hydrogel character of the material was irreversibly lost due to the covalent capture reaction. In contrast to the thin fibers observed in supramolecular F*Dopa*, we observed a bundling of the fibrous assemblies into thick fibers by Cryo-EM (Figures 6a/e,). Due to insolubility from covalent capture, we utilized scanning electron microscopy (SEM) to successfully confirm the presence of fibrous structures at both pH values using previously established methods (Figures 6b/f ). ^63^

**Fig. 6.**
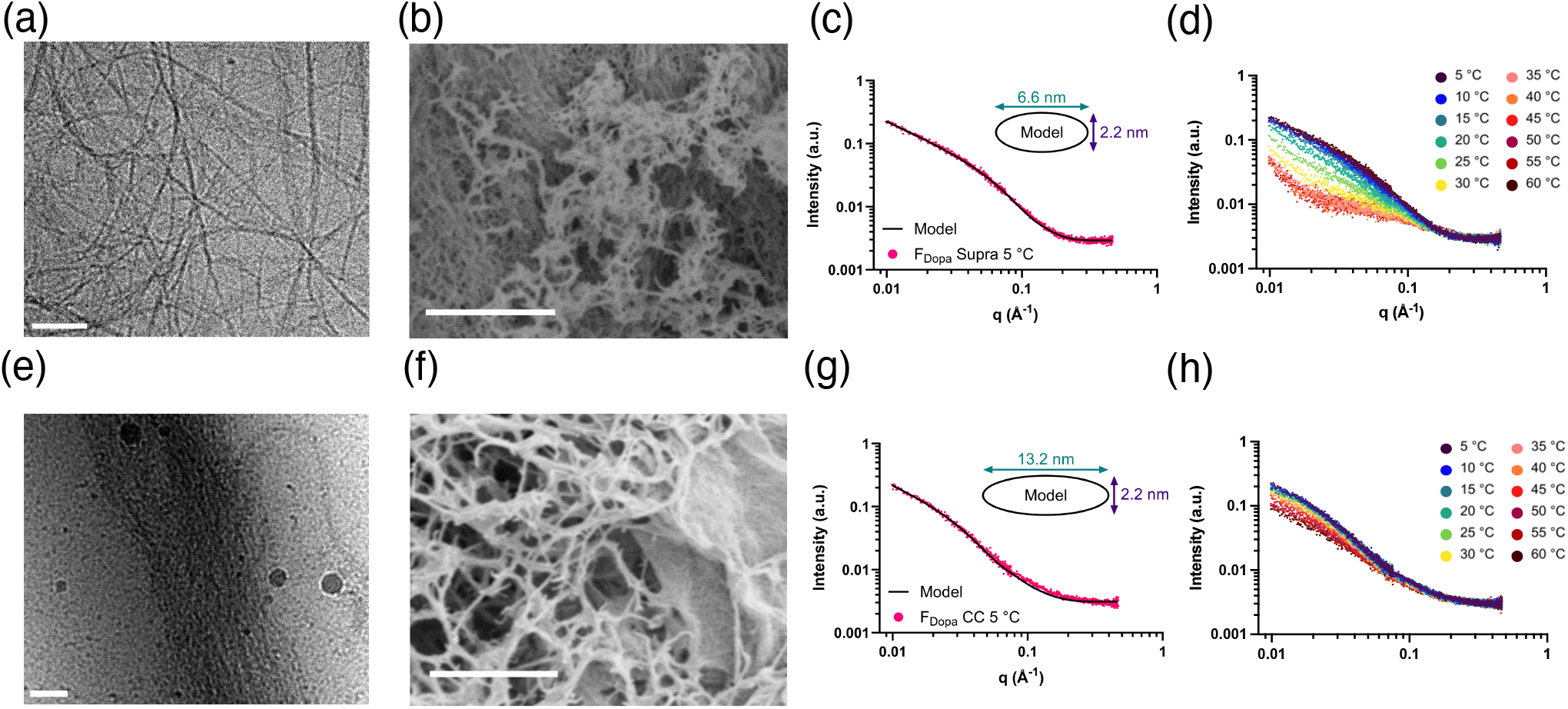
Covalently stabilizing fibrous collagen mimetic assemblies through Lys-Dopa interactions. (a) Cryogenic electron microscopy (Cryo-EM) of supramolecular assemblies of F_*Dopa*_ at pH = 4.5, the scale bar is 100 nm. (b) Scanning electron microscopy (SEM) of F_*Dopa*_ at pH = 4.5 also shows a fibrous composition; the scale bar is 1 *µ*m. (c) Scattering curve of supramolecular F_*Dopa*_ fibers at 5 °C and pH = 4.5 modeled as 400 nm long elliptical cylinder scattering objects. (d) SAXS curves of the supramolecular fibers from 5-60 °C show a decrease in fiber signal with increasing temperature until the fibers are completely disassembled at ca. 40 °C. (e) Cryo-EM of covalently captured fibrous assemblies of F_*Dopa*_ at pH = 9.5, the scale bar is 100 nm. (f) SEM of F_*Dopa*_ at pH = 9.5 retains a fibrous composition, and the scale bar is 1 *µ*m. (g) Scattering curve of covalently captured F_*Dopa*_ fibers at 5 °C and pH = 9.5 modeled as 400 nm long elliptical cylinder scattering objects showing that the major diameter doubles compared to the supramolecular fibers. (h) Scattering curves of covalently captured F_*Dopa*_ fibers show only a minimal decrease in fiber signal from 5-60 °C.

To measure the thermal stabilization of the covalently captured F*Dopa* fibers, we performed small-angle X-ray scattering (SAXS) experiments from 5-60 °C on both the supramolecular and covalently captured fibers. SAXS is able to report on the average macromolecular structure of the fibers in situ as opposed to Cryo-EM, which requires samples to be frozen, and CD, which can only be used to investigate the secondary structure and cannot report on the presence of fibers directly.

Scattering curves of both the supramolecular and covalently captured samples at 5 °C confirmed that both formed fibers that could be modeled as elliptical cylinders (Figures 6c/g and Table S2). Both the supramolecular and covalently captured fibers have a minor diameter of 2.2 nm, but the covalently captured system had an average major diameter that was 2-fold larger (6.6 vs 13.2 nm). These results are consistent with Cryo-EM, which shows that the fibers have a higher propensity to bundle together upon being covalently captured. These samples were then gradually heated to 60 °C, and SAXS measurements were taken every 5 °C to monitor changes in the fiber structure. Upon heating, the supramolecular fibers started to disappear, as seen as a temperature-dependent decrease in the scattering intensity (Figure 6d). By 40 °C, the scattering curve was consistent with the presence of disordered peptide, suggesting that the supramolecular fibers were completly disassembled at this temperature. The covalently captured fibers, in contrast, were much more resistant to heat, and we observed only a modest decrease in scattering intensity over the entire temperature range (Figure 6h).

The average scattering intensity plotted as a function of temperature shows that the supramolecular fibers have a *Tm* of 25 °C, as determined by finding the inflection point of the sigmoidal curve fit (Figure S17a). The *Tm* for the covalently captured system could not be accurately calculated because the fibers did not completely disassemble over the investigated temperature range. The thermal stability of the supramolecular F*Dopa* fibers was further investigated by conducting a CD temperature sweep, which resulted in a *Tm* of 18 °C (Figure S17b). Taken together, these data suggest that the Lys – Dopa covalent capture scheme can stabilize higher-order CMP fiber assemblies.

## Conclusion

Crosslinking chemistry is an essential post-translational modification in natural collagen. Synthetic crosslinking of collagen mimetic peptides has been previously achieved with disulfide bonding, isopeptide amidation. Through simple pH adjustment, this work demonstrates a facile reagent-free method to covalent capture triple helices through oxidative crosslinking between Dopa and Lys. The axial geometry was preferential in creating a covalent modification. While the reaction did not lead to discrete triple helical purification like in the case of amidation, it was demonstrated that the reaction could stabilize simple homotrimers and complex heterotrimers with an abundance of other reactive Lys side chains. Finally, the covalent stabilization of collagen mimetic peptide fibers was achieved. It may be a synthetic strategy for generating novel collagen mimetic biomaterials that could be used to mimic the extracellular matrix. Lastly, supported by the expression of Dopa and Lys in proximity to each other in mussels, ^68^ the presence of both Lys and Dopa residues in human collagen, ^13^ and the formation of lysyl tyrosylquinone (LTQ) crosslinking in lysine oxidase, ^15^ we hypothesize this could be a reasonable covalent crosslinking stabilizing natural collagens.

## Data availability

The datasets supporting this article have been uploaded as part of the SI.

## 4 Author Contributions

Carson C. Cole: conceptualization, methodology, investigation, writing. Brett H. Pogostin: conceptualization, methodology, investigation, writing. Vardan H. Vardanyan: investigation, data curation. Kiana Cahue: conceptualization, investigation, data curation. Thi H. Bui: investigation, data curation. Adam C. Farsheed: investigation, data curation. Joseph W.R. Swain: investigation, data curation. Jonathan Makhoul: investigation, data curation. Marija Dubackic: investigation, data curation. Peter Holmqvist: investigation, data curation. Ulf Olsson: conceptualization, funding acquisition. Anatoly B. Kolomeisky: supervision, funding acquisition. Kevin J. McHugh: editing, writing, supervision, funding acquisition. Jeffrey D. Hartgerink: editing, writing, supervision, funding acquisition.

## 6 Acknowledgements

The authors would like to acknowledge Crispin Hetherington and L. Tracy Yu for their technical assistance and insights. This work was funded in part by the National Science Foundation (CHE 2203937), the National Science Foundation Graduate Research Fellowship (Grant No. 1842494), the Welch Foundation (C-2141) and the Swedish Research Council (2020-04633). This work was partly supported by the Big-Data Private-Cloud Research Cyberinfrastructure MRI award funded by the NSF under grant CNS-1338099 and by Rice University’s Center for Research Computing (CRC). This work benefited from using the SasView application, originally developed under NSF award DMR-0520547. SasView contains code developed with funding from the European Union’s Horizon 2020 research and innovation program under the SINE2020 project, grant agreement No 654000.

## 1 Methods Continued

### 1.1 UV-Vis Characterization

Homotrimer and heterotrimer samples were diluted to 0.15 mM for total peptide concentration and in both cases of supramolecular and covalently captured peptides. Upon dilution, the absorbance measurements were carried out in a quartz cuvette on a UV-Vis Spectrophotometer Evolution 220 (Thermo Scientific, USA).

### 1.2 ATR-FTIR

Attenuated Total Reflectance Fourier Transform Infrared spectroscopy (ATR-FTIR) was used to analyze the secondary structure of DOPA-containing peptides and controls using a NicoletTM iS20 FTIR spectrometer (Thermo Fisher Scientific, Waltham, MA). Each sample was dried on the window with a stream of nitrogen gas and measured using 30 scans at a resolution of 4 cm^−1^. Intensities were normalized to the highest intensity of each sample.

### 1.3 Rheology

Rheology was performed on an AR-G2 rheometer (TA Instruments, Delaware, USA) with a 12 mm parallel plate. 100 *µ*L of peptide was extruded onto the stage, and the parallel plate was lowered to a gap of 550 *µ*m. Any excess peptide solution was scraped away, and 3 drops of mineral oil were applied surrounding the plate to prevent evaporation during testing. The plate was lowered to a final gap of 500 *µ*m. The following tests were performed at the accompanying parameters: Strain sweep: 0.1 to 100% strain at 1 rad s^−1^. Shear sweep: 0.01 to 100 s^−1^.

## 2 Peptide Characterization

**Figure S1:**
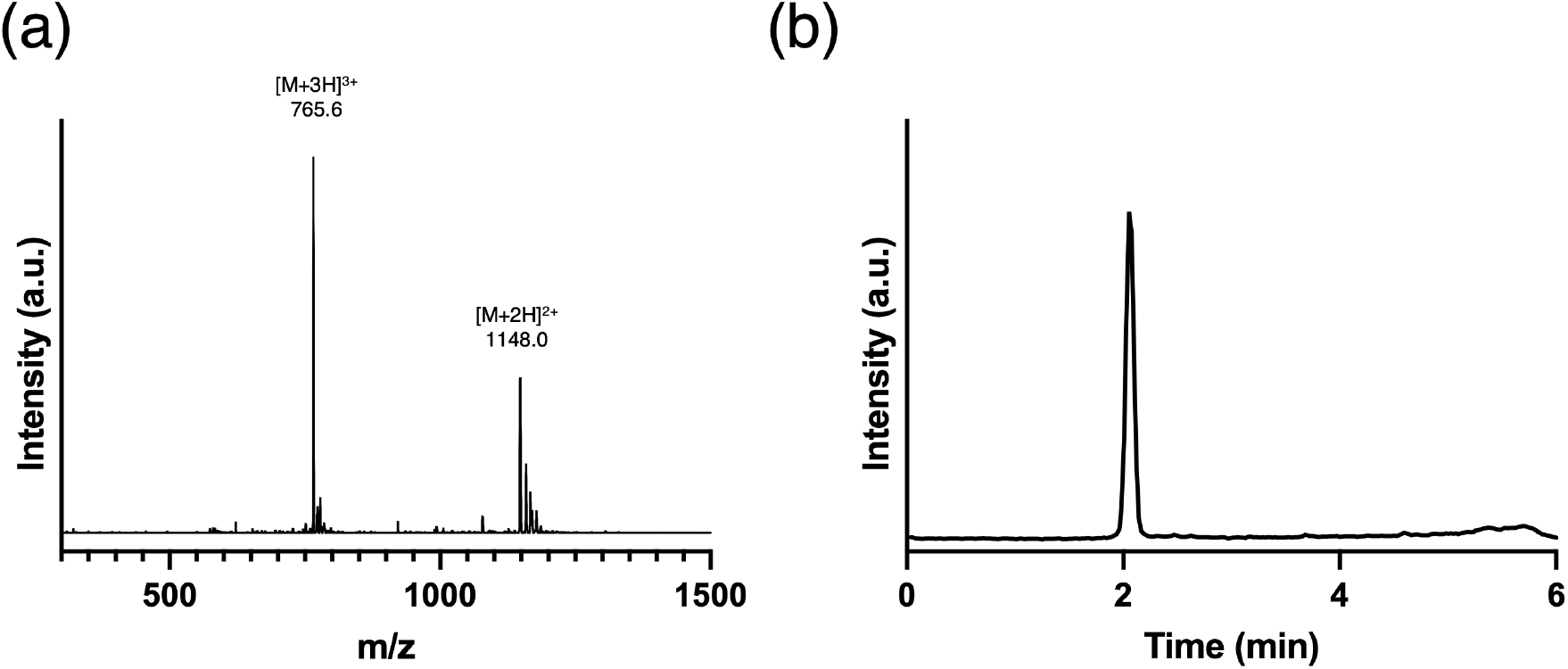
KGDopaO Characterization: a) mass spectra (Expected: [M + H]^+^ = 2295.4; Observed: 2295.0). b) Pure liquid chromatography trace.

**Figure S2:**
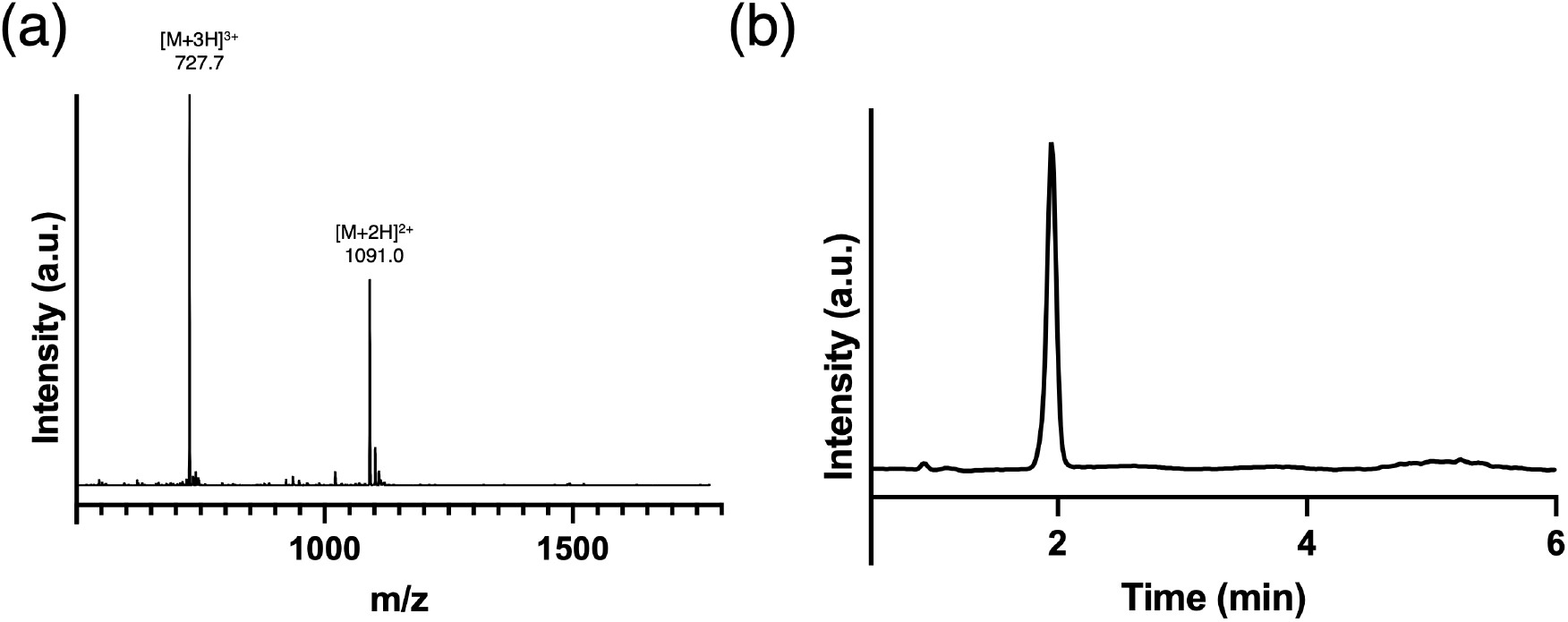
KGDopaO Characterization with a 6,9 glycine deletion: a) mass spectra (Expected: [M + H]^+^ = 2181.3; Observed: 2181.0). b) Pure liquid chromatography trace.

**Figure S3:**
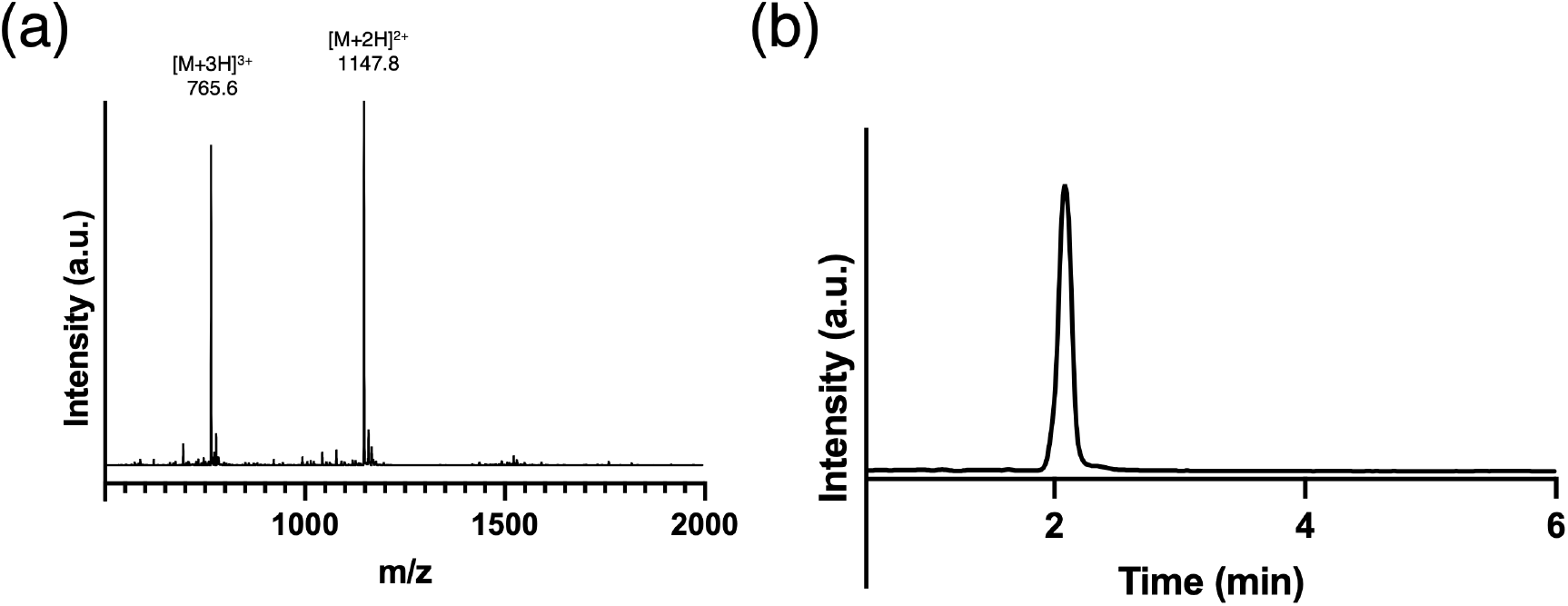
OGDopaK Characterization: a) mass spectra (Expected: [M + H]^+^ = 2295.4; Observed: 2294.6). b) Pure liquid chromatography trace.

**Figure S4:**
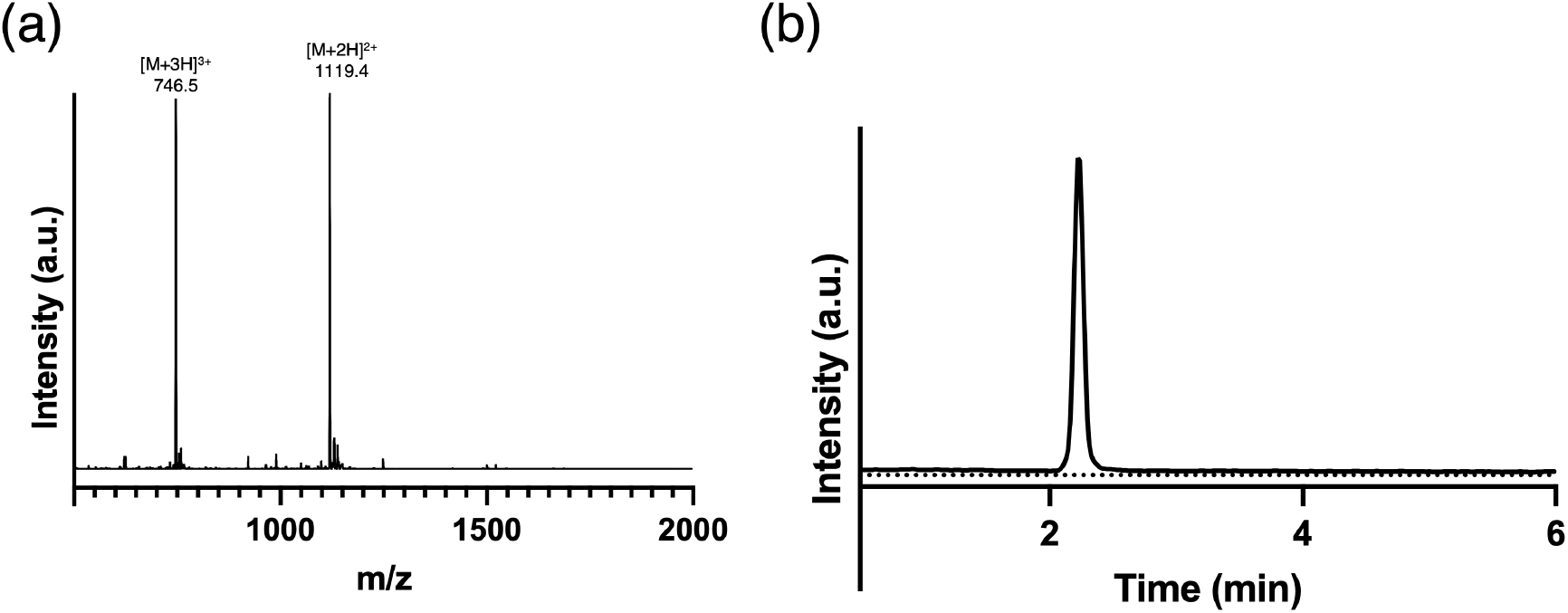
AGDopaO Characterization: a) mass spectra (Expected: [M + H]^+^ = 2238.2; Observed: 2237.8). b) Pure liquid chromatography trace.

**Figure S5:**
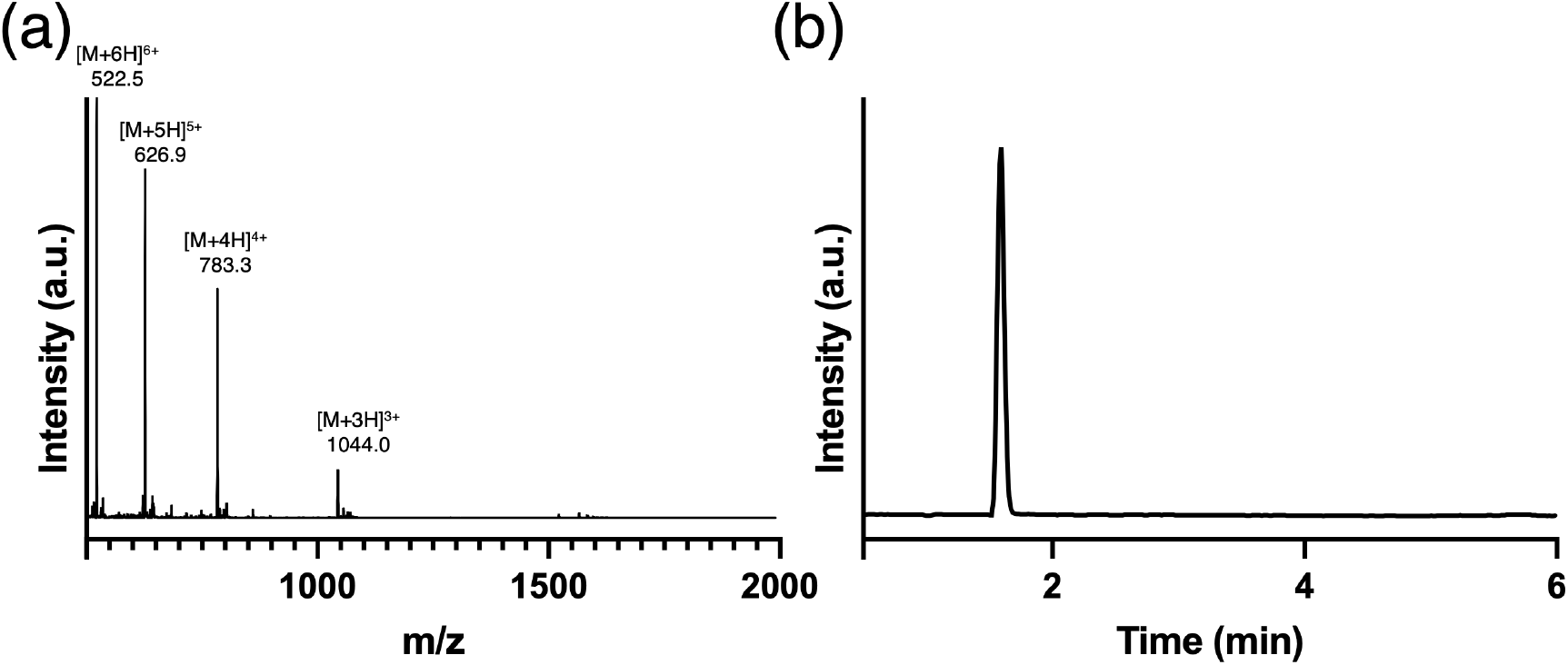
HT3A Characterization: a) mass spectra (Expected: [M + H]^+^ = 3128.7; Observed: 3130.0). b) Pure liquid chromatography trace.

**Figure S6:**
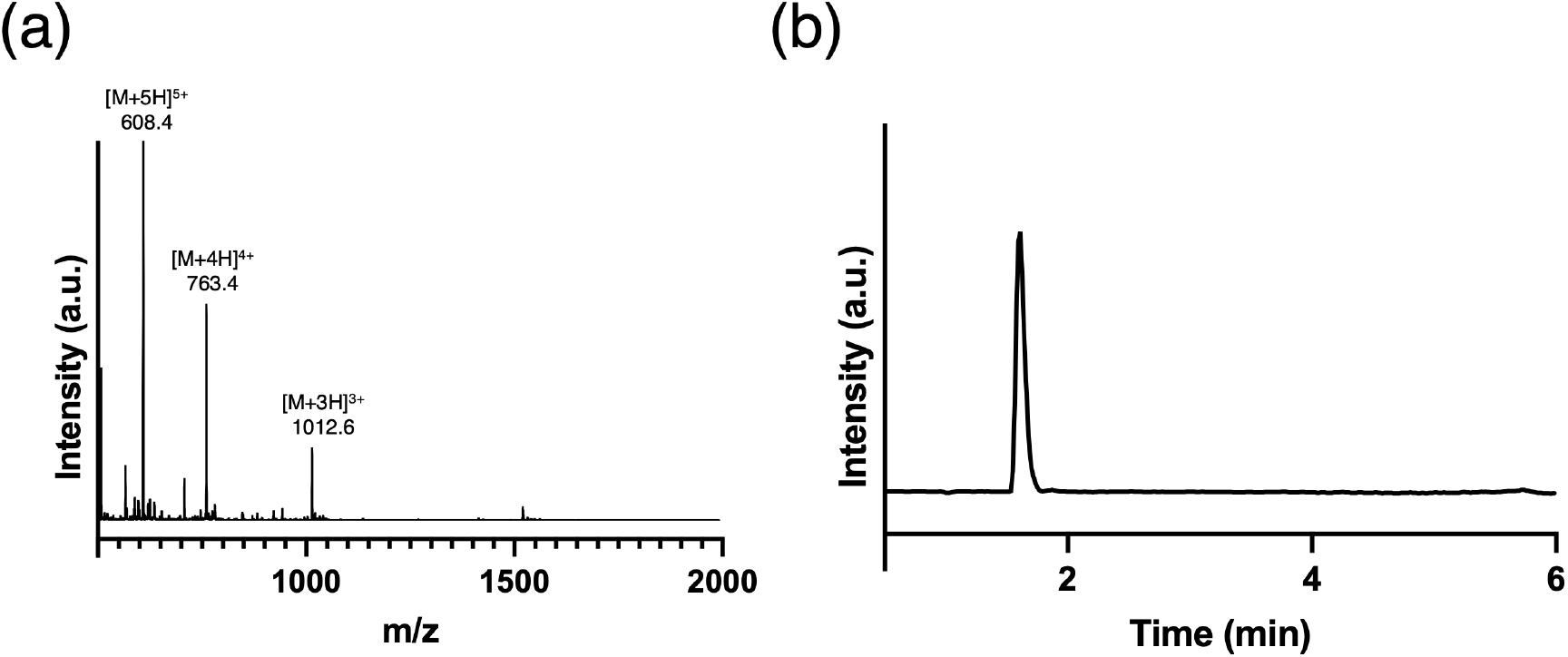
HT3B Characterization: a) mass spectra (Expected: [M + H]^+^ = 3037.5; Observed: 3037.8) b) Pure liquid chromatography trace.

**Figure S7:**
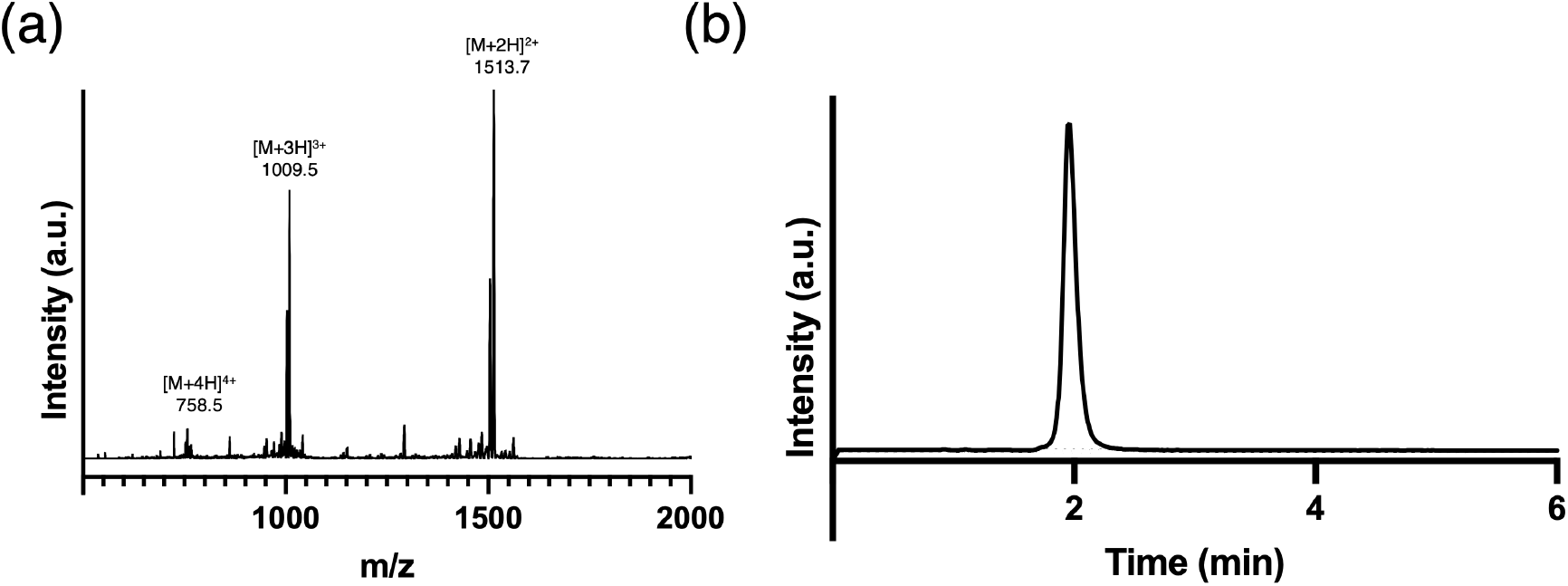
HT3C-22-Dopa Characterization: a) mass spectra (Expected: [M + H]^+^ = 3024.1; Observed: 3025.7). b) Pure liquid chromatography trace.

**Figure S8:**
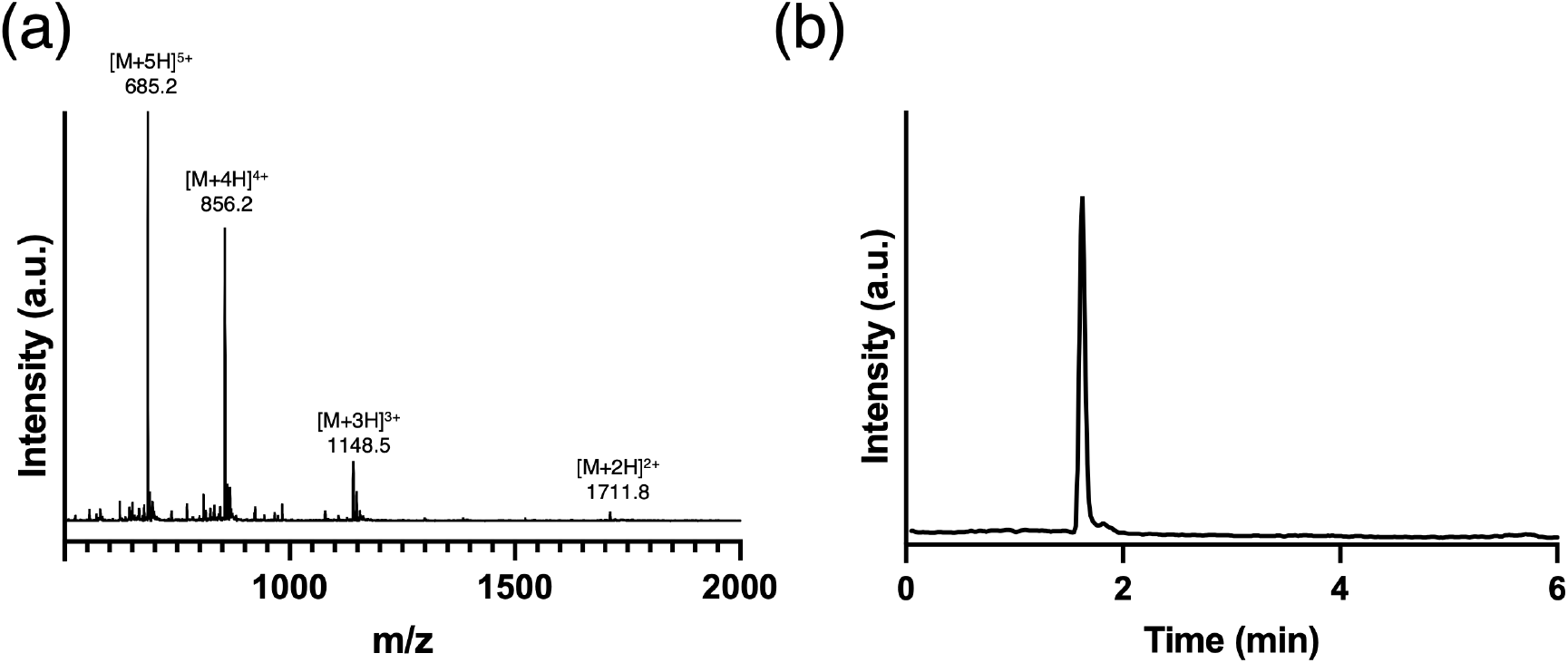
F_*Dopa*_ Characterization: a) mass spectra (Expected: [M + H]^+^ = 3422.6; Observed: 3421.6) b) Pure liquid chromatography trace.

## 3 Homotrimer Characterization

**Figure S9:**
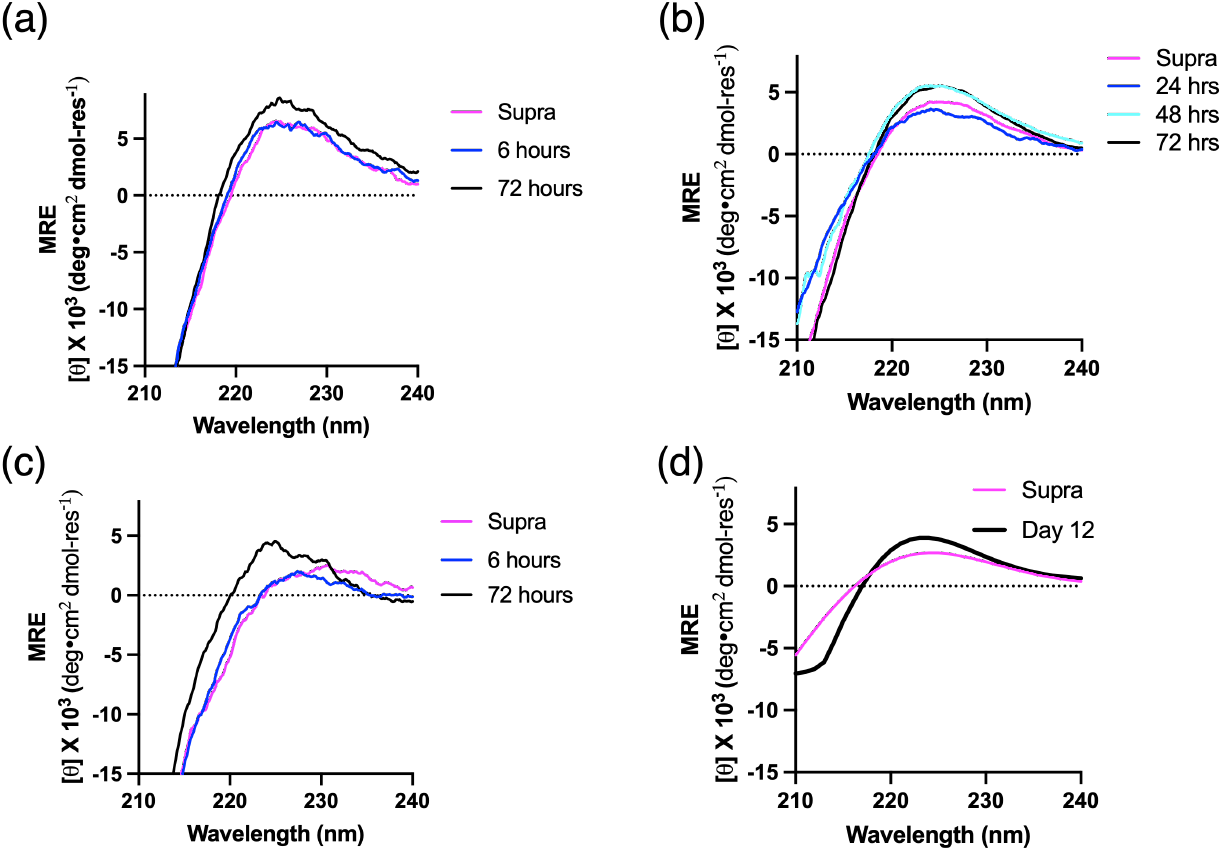
Circular dichroism spectra of homotrimers and F_*Dopa*_. (a) KGDopaO CD spectra of supramolecular (supra) and covalently captured species. (b) OGDopaK CD spectra of supra and covalently captured species. (c) AGDopaO CD spectra of supra and covalently captured species. (d) F_*Dopa*_ CD spectra of supra and covalently captured. Supra spectra collected in 1 mM bicarbonate buffer at pH 4.5. Covalently captured spectra collected in 1 mM bicarbonate buffer (pH 9.5) and at 5 °C.

**Figure S10:**
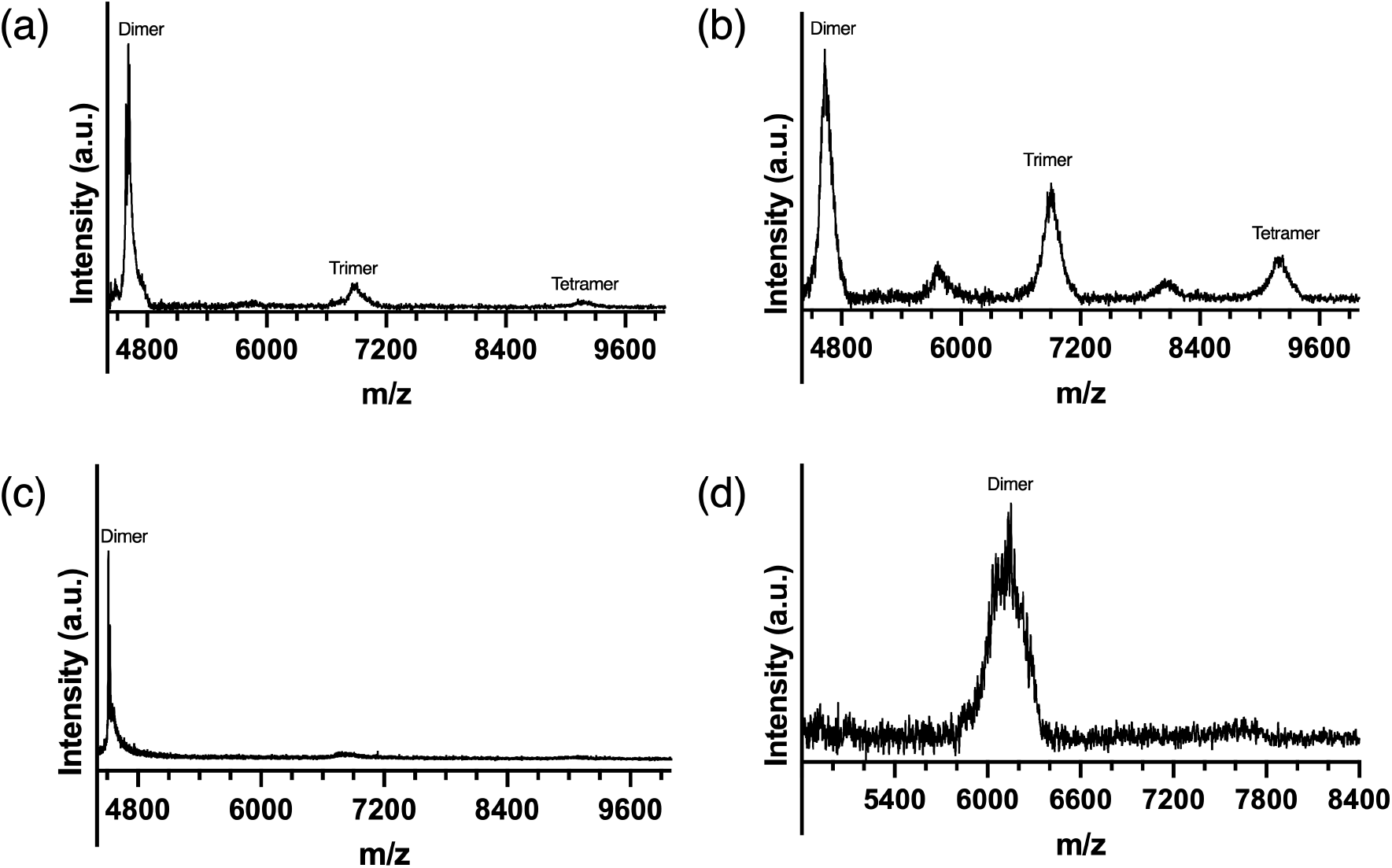
Higher order characterization of homotrimers with MALDI ToF MS in linear positive mode. (a) KGDopaO spectra shows clear dimer (expected 4590.9 m/z), trimer (expected 6886.3 m/z) and tetramer (expected 9181.7 m/z) formation. (b) OGDopaK spectra shows clear dimer (expected 4590.9 m/z), trimer (expected 6886.3 m/z), tetramer (expected 9181.7 m/z) and possible +2 ionization states of higher-order species (e.g., pentameric, heptameric). (c) AGDopaO spectra shows dimer formation (expected 4475.6 m/z). (d) ABC-Dopa shows a mass correlating to the dimer formation within the range of the B – C dimer (expected 6061 m/z).

**Figure S11:**
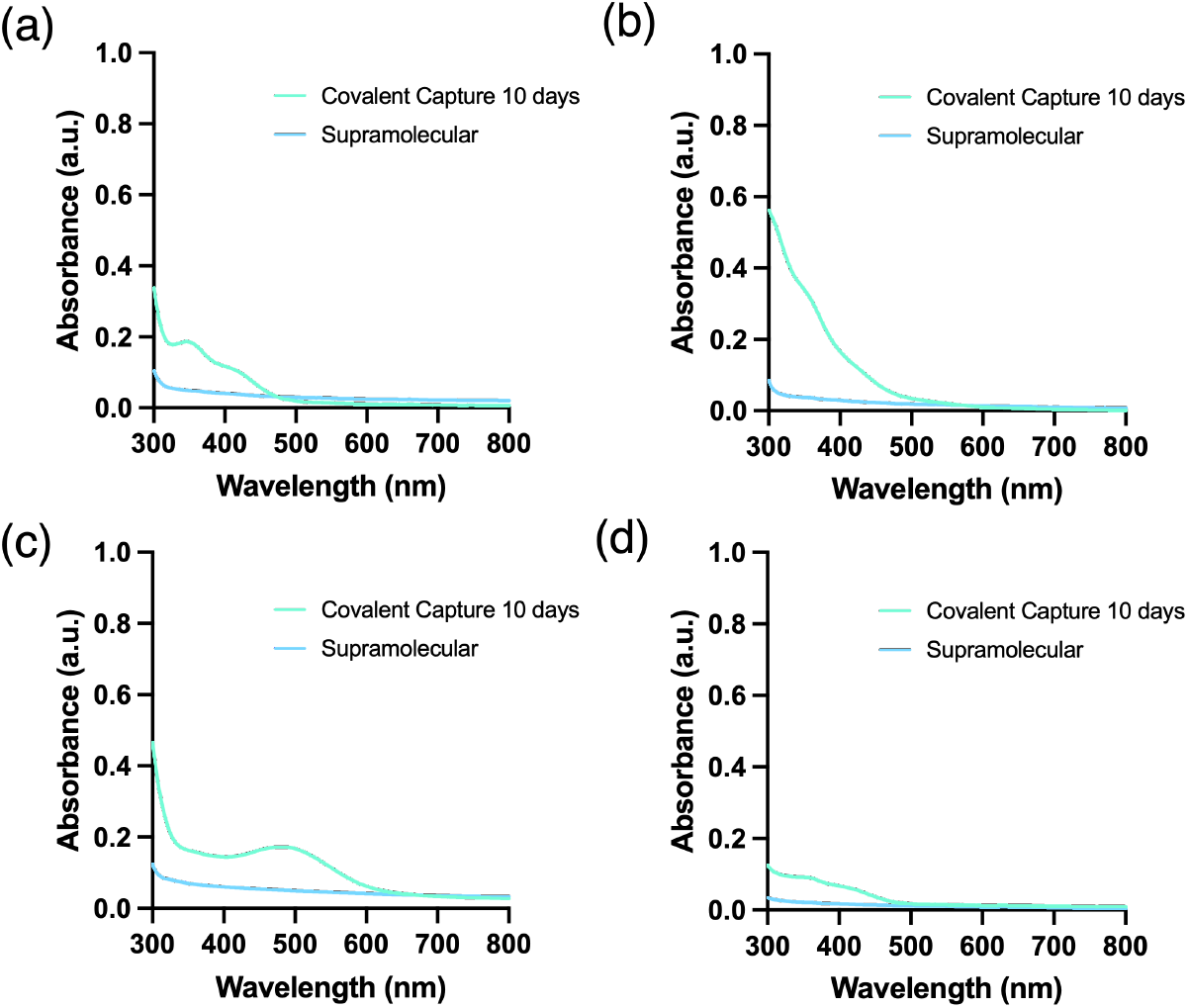
UV-Vis spectra of homotrimers and a ABC-type Heterotrimer. (a) KGDopaO spectra after reacting at pH = 9.5 compared to the supramolecular homotrimer (pH = 4.5). (b) OGDopaK spectra at pH = 9.5 compared to the supramolecular homotrimer (pH = 4.5). (c) AGDopaO spectra at pH = 9.5 compared to the supramolecular homotrimer (pH = 4.5). A distinct peak at ca. 495 nm correlates to the distinct red color. (d) ABC-22 Dopa spectra at pH = 4.5 and 9.5.

**Figure S12:**
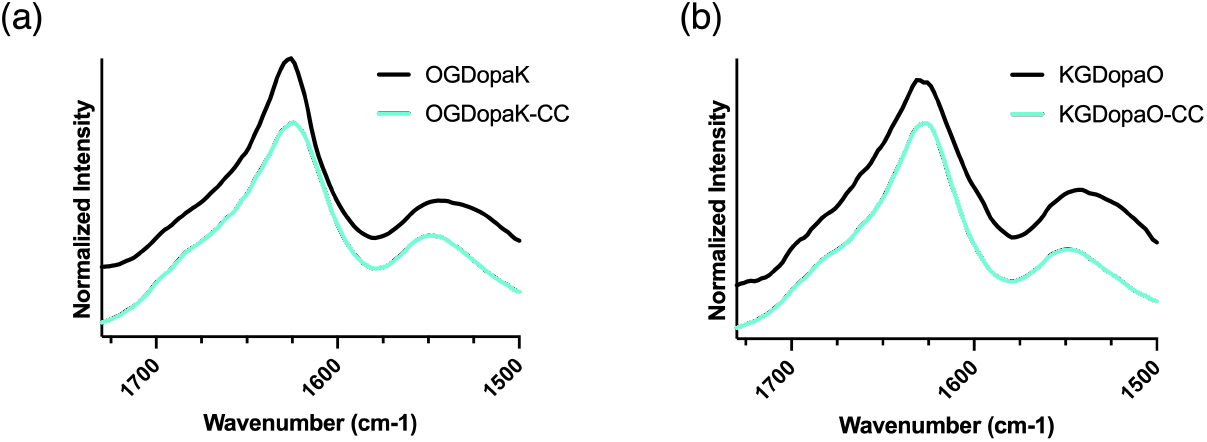
ATR-FTIR of OGDopaK and KGDopaO. (a) IR spectra of OGDopaK supra and covalently captured (CC) samples show little shift in peak maxima (b) IR spectra of KGDopaO supra and CC samples show no shift in peak maxima, supporting a triple helical tertiary structure in both cases.^1^

**Figure S13:**
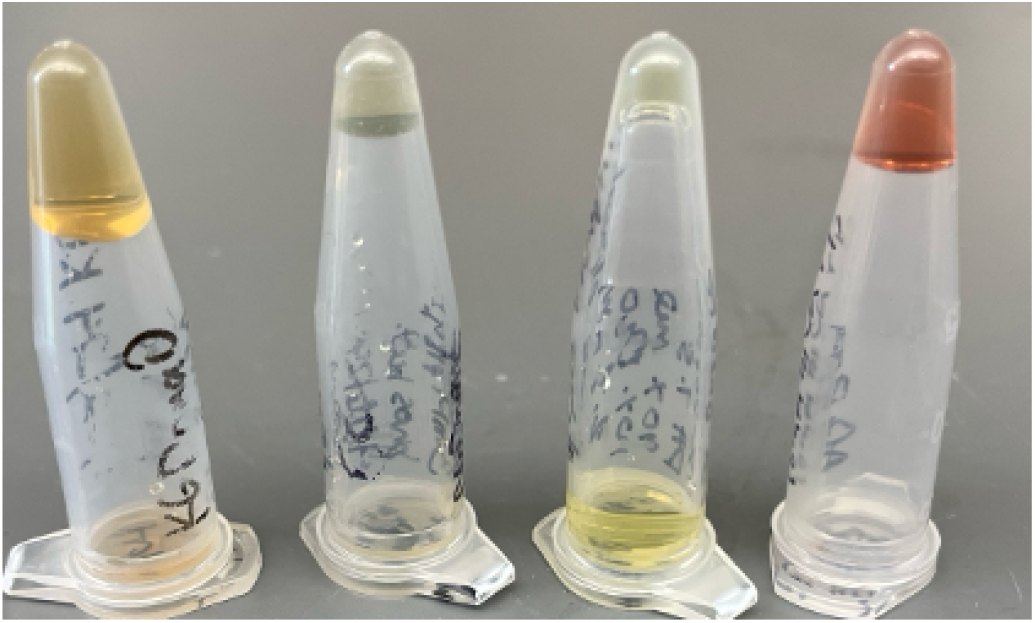
Eppendorf tubes of homotrimer and heterotrimer solutions. From left to right: KGDopaO, OGDopaK, ABC-Dopa (pH = 7.4) and AGDopaO at 3mM and pH = 9.5.

**Figure S14:**
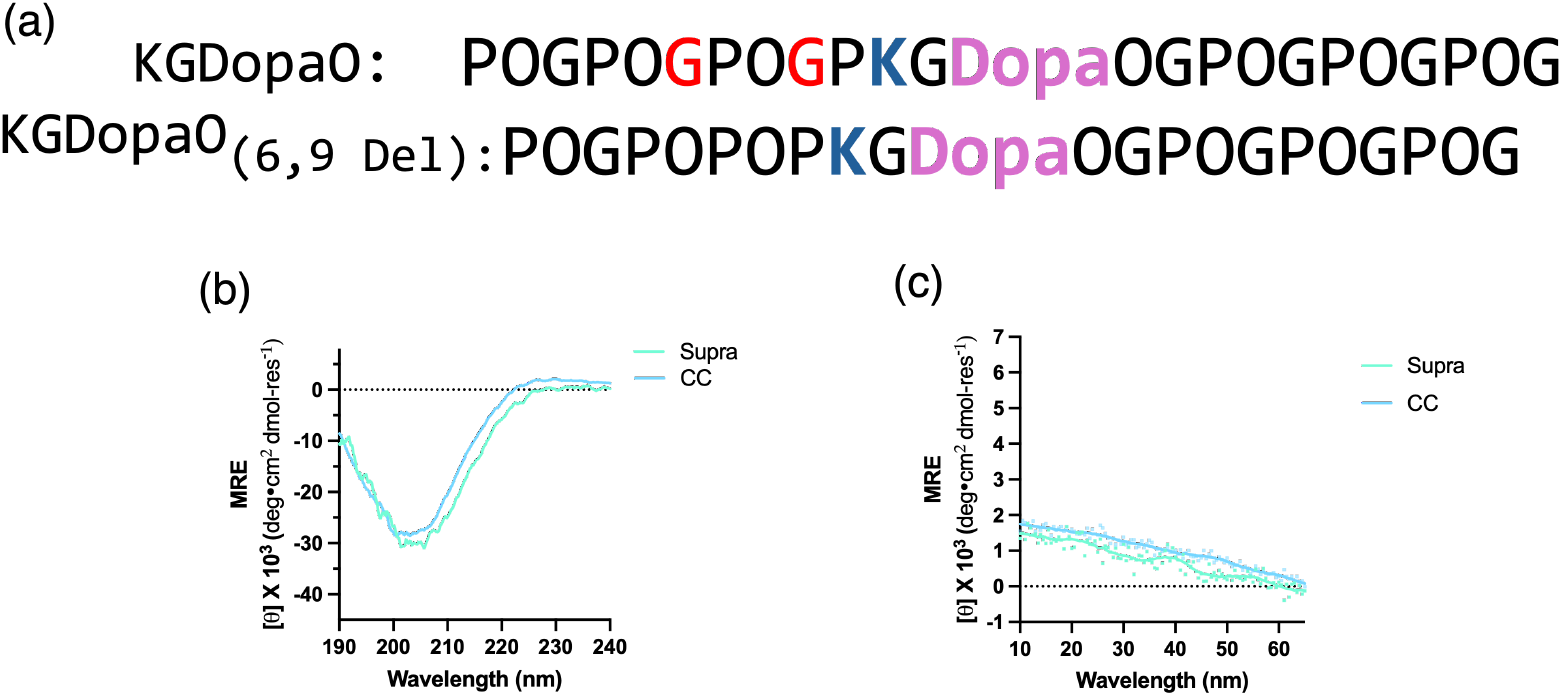
KGDopaO 6,9 Gly deletion characterization with circular dichroism. (a) Sequence comparison of KGDopaO and KGDopaO with glycine deletions at positions 6 and 9. Deleted glycines are highlighted in red in the original sequence of KGDopaO. (b) CD spectra of covalently captured and supramolecular KGDopaO 6,9 Gly deletion. (c) CD thermal melts of covalently captured and supramolecular KGDopaO 6,9 Gly deletion. There is a lack of strong thermal transition for both samples. All samples were covalently captured at pH = 9.5 and recorded after twenty-five days.

**Figure S15:**
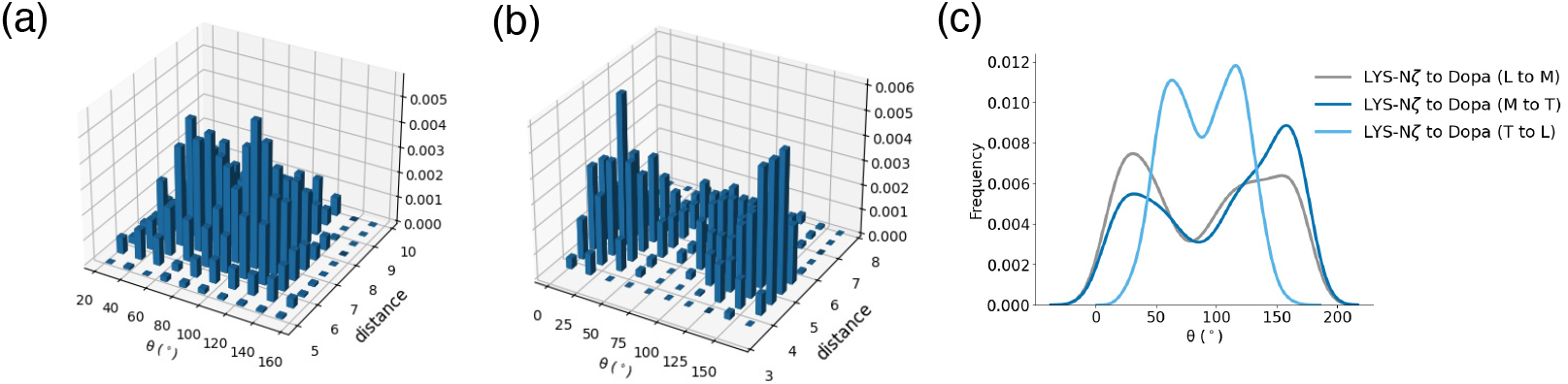
Angle characterization of cation-*π* interactions in KGDopaO. (a) Lateral cation-*π* interaction with distance and angle correlations. (b) Axial cation-*π* interaction. (c) Axial versus lateral interaction angle population curves

## 4 Fibers Characterization

**Figure S16:**
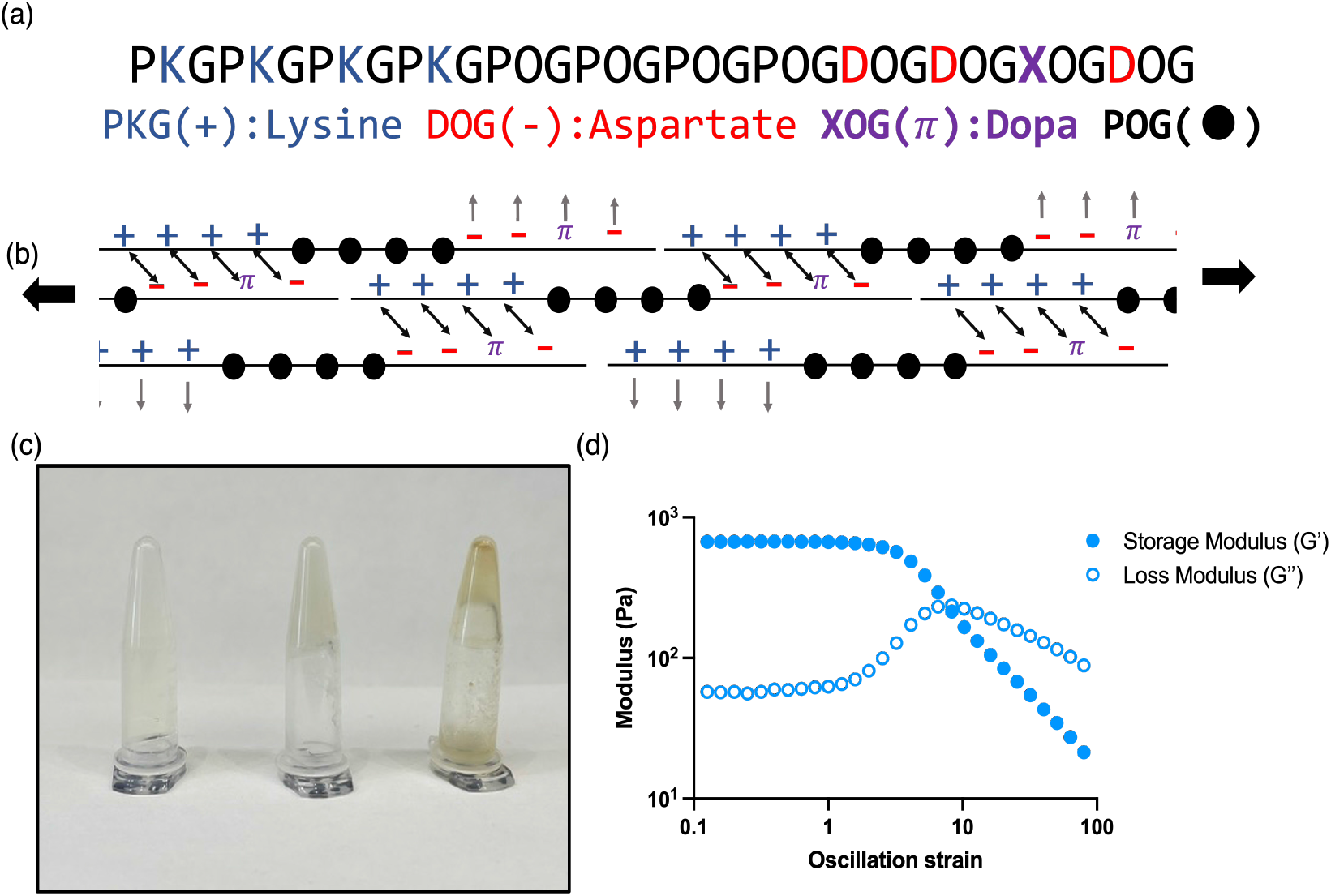
Sequence design and rheological characterization of F_*Dopa*_. (a) Sequence of polymerizing F_*Dopa*_. (b) Proposed polymerization mechanism of a sticky-ended peptide. (c) F_*Dopa*_ inverted eppendorf tubes at pH 4.5, 7.4 and 9.0. (d) Rheology of pH 4.5 hydrogel of F*Dopa*.

**Figure S17:**
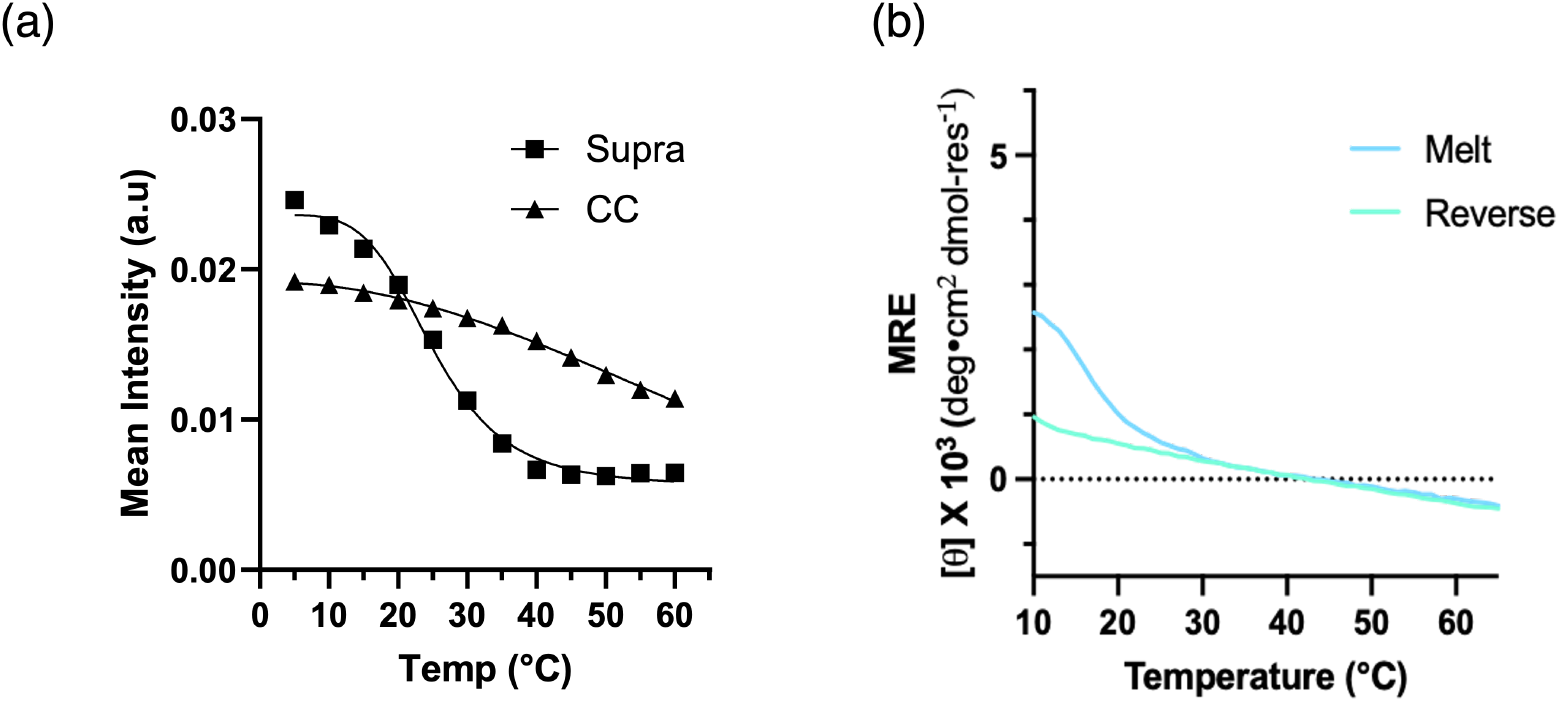
Scattering and CD melt comparison of F_*Dopa*_. (a) Scattering melting curves taken by plotting the average intensity from Figure 6d/h. Supra is F_*Dopa*_ at pH 4.5, and covalent capture (CC) is F_*Dopa*_ at pH 9.5. (b) F_*Dopa*_ at pH 4.5 characterized with CD demonstrates that PPII signal is not directly related to fiber stability. The observed hysteresis is typical for a supramolecular reverse melt. After covalent capture at pH 9.5, the precipitate made CD characterization not possible.

**Table S1:** Reaction Conditions for Covalent Capture

| Sample | Instrument | Buffer | pH | Reaction Temp. (°C) | Conc. [mM] |
| --- | --- | --- | --- | --- | --- |
| KGDopaO | CD | 10 mM NaHCO<sub>3</sub> | 9.5 | 4.0 | 3.00 |
| OGDopaK | CD | 10 mM NaHCO<sub>3</sub> | 9.5 | 4.0 | 3.00 |
| AGDopaO | CD | 10 mM NaHCO<sub>3</sub> | 9.5 | 4.0 | 3.00 |
| KGDopaO<sub>6,9GlyDeletion</sub> | CD | 10 mM NaHCO<sub>3</sub> | 9.5 | 4.0 | 3.00 |
| ABC Het. | CD | 10 mM PB | 7.4 | 30.0 | 3.00 |
| ABC Het. | NMR | 9 mM PB/10% D<sub>2</sub>O | 7.4 | 30.0 | 2.70 |
| $F_{Dopa}$ | CD/SAXS | None | 9.5 | 4.0 | 3.00 |

**Table S2:** Fitting parameters for SAXS fits of F_*Dopa*_

| Sample | $F_{Dopa}$ Supra | $F_{Dopa}$ CC |
| --- | --- | --- |
| Scale | $6 \times 10^{-4}$ | $3 \times 10^{-4}$ |
| Background | $2.9 \times 10^{-3}$ | $2.9 \times 10^{-3}$ |
| Minor Radius (Å) | 11 | 11 |
| axis ratio | 3 | 6 |
| length (nm) | $4 \times 10^2$ | $4 \times 10^2$ |
| SLD Peptide (Å<sup>-2</sup>) | $1.27 \times 10^{-5}$ | $1.27 \times 10^{-5}$ |
| SLD Solvent (Å<sup>-2</sup>) | $9.44 \times 10^{-6}$ | $9.44 \times 10^{-6}$ |

